# Conserved transcriptional plasticity, not local adaptation, dominates early climate responses across wild and cultivated apples

**DOI:** 10.64898/2026.01.15.698562

**Authors:** Ronan Dadole, Xilong Chen, Jorge Sandoval, Anthony Venon, Maxime Criado, Noemie Delpouve, Carine Remoué, Simon Chollet, Amandine Hansart, Francois Laurens, Laurence Feugey, Thomas Kirisits, Tudor M. Ursu, Anamaria Roman, Gayle M. Volk, Georgina Alins, Natalia Conde e Silva, Komlan Avia, Stephane Bazot, Amandine M. Cornille

## Abstract

Perennial crops respond to climate change through phenotypic plasticity and local adaptation, yet how these responses are orchestrated at the molecular level and reshaped by domestication remains poorly understood.

We grew 1,249 seedlings from five populations of M. domestica, M. orientalis, and M. sylvestris under four simulated European climates, quantified 12 phenotypic traits, performed RNA sequencing on 96 individuals, and integrated these data with previously published genome-wide polymorphism data from M. sylvestris to disentangle conserved climate responses from local adaptation. Climate was the dominant driver of phenotypic and transcriptional variation, revealing 344 conserved climate-responsive genes enriched for stress responses, nutrient metabolism, and cell wall biosynthesis. These genes are under strong purifying selection and carry fewer deleterious mutations. In contrast, genome-environment association analyses identified 217 loci associated with temperature and precipitation gradients, only a subset of which showed climate-responsive expression and elevated genomic differentiation. Population-level expression divergence closely mirrored neutral genetic differentiation, indicating that most transcriptomic divergence reflects demographic history rather than adaptive regulatory evolution. Domestication reshaped molecular diversity and mutation load without disrupting the conserved climate-response program.

Climate responses in apple are primarily mediated by an evolutionarily conserved transcriptional program maintained by strong purifying selection, whereas local adaptation relies on largely distinct, population-specific loci. These findings reveal that conserved plasticity constitutes the dominant molecular response to climatic variation, whereas local adaptation contributes a secondary, population-specific layer of evolutionary change.

## Introduction

Rapid anthropogenic climate change is placing unprecedented pressure on both natural and cultivated plant populations (Jump and Peñuelas 2005). While past climatic shifts, such as the Pleistocene glacial-interglacial cycles, shaped species distributions over millennia, contemporary warming is occurring at a hitherto unknown pace (Hewitt 1999; Hewitt 2004; Williams et al. 2008; Corlett and Westcott 2013). Whether perennial species can adapt fast enough to survive this acceleration is a critical question in evolutionary ecology, with implications for biodiversity conservation, food security, and a wide range of other ecosystem services (Thomas et al. 2004; Rellstab et al. 2021; Capblancq et al. 2023; Rezaei et al. 2023; Cortés 2024; Alseekh et al. 2025).

Plants respond to environmental change through local adaptation, whereby genetic differentiation increases fitness in native environments, and through phenotypic plasticity, whereby a single genotype produces different phenotypes across environments (Kudoh 2016; Fox et al. 2019; Anderson and Song 2020; Satake et al. 2022). Although common-garden and reciprocal transplant experiments have documented extensive phenotypic variation (Savolainen et al. 2013; de Villemereuil et al. 2016; Sork 2018; Benito Garzón et al. 2019; Schwinning et al. 2022), the molecular mechanisms underlying these responses remain poorly understood. Disentangling plasticity from local adaptation is particularly challenging because population history can generate signals that resemble adaptation if not properly accounted for.

Long-lived perennial trees provide an exceptional system for addressing these questions. Their long lifespan requires them to buffer short-term environmental fluctuations while maintaining fitness over decades, implying a complex interplay between phenotypic plasticity and adaptive differentiation (Hampe and Petit 2005; Kremer et al. 2012; Lobo et al. 2018; Honjo and Kudoh 2019; Anderson et al. 2021; Kremer et al. 2025). Recent advances in genomics now enable integrated analyses of phenotypes, gene expression, and genome-environment associations, providing unprecedented opportunities to uncover the molecular basis of climate responses in natural populations (Capblancq et al. 2023; Kremer et al. 2025; Verrico et al. 2025). In perennial crops, domestication adds an additional evolutionary dimension by reshaping demographic history, regulatory networks, and selective regimes. These processes can modify gene expression, alter the efficiency of purifying selection, and influence the accumulation of deleterious variation, yet whether they constrain or preserve climate-responsive pathways remains largely unknown (Gaut et al. 2015; Swinnen et al. 2016; Bosse et al. 2019; Andersson and Purugganan 2022; Gao et al. 2023).

Apple (*Malus domestica Borkh.*) and its wild relatives provide an ideal model to investigate these questions because their domestication history and population structure are exceptionally well characterized. The cultivated apple originated primarily from the Central Asian wild apple, *M. sieversii*, with additional contributions from *M. orientalis* and *M. sylvestris* during its spread into Europe (Cornille et al. 2012; Cornille et al. 2014). Wild populations occupy contrasting climatic environments and retain clear signatures of historical demographic processes and local differentiation (Cornille et al. 2015; Chen et al. 2023), making this system particularly well-suited to disentangling conserved climate-responsive mechanisms from population-specific adaptation.

Here, we leverage the well-characterized domestication histories and population genetic structures of wild and cultivated apples (Cornille et al. 2012; Cornille et al. 2013; Cornille et al. 2014; Cornille et al. 2015; Cornille et al. 2019; Chen et al. 2023; Chen et al. 2026) to investigate how phenotypic plasticity, local adaptation, and domestication shape climate responses in *M. domestica, M. orientalis, and M. sylvestris*. Specifically, we ask whether climate responses are primarily driven by conserved plasticity or population-specific adaptation, whether they involve a shared transcriptional program across lineages, and how domestication has reshaped the genomic context in which these responses evolve. To address these questions, we grew seedlings from five apple populations under four simulated European climate regimes in climate-controlled ecotrons (Fig. 1a,b). We quantified survival and twelve morphophysiological traits and integrated these data with transcriptome sequencing, population genomic analyses, and genotype-environment association analyses (Fig. 1a; supplementary fig. S1). Together, these complementary datasets revealed a conserved transcriptional program underlying early climate responses across wild and cultivated apples, whereas signatures of local adaptation were comparatively limited and largely restricted to wild populations. Domestication reshaped patterns of gene expression and genome evolution without fundamentally altering this conserved climate-responsive program, providing a framework for understanding and predicting climate resilience in perennial crops.

**Figure 1.**
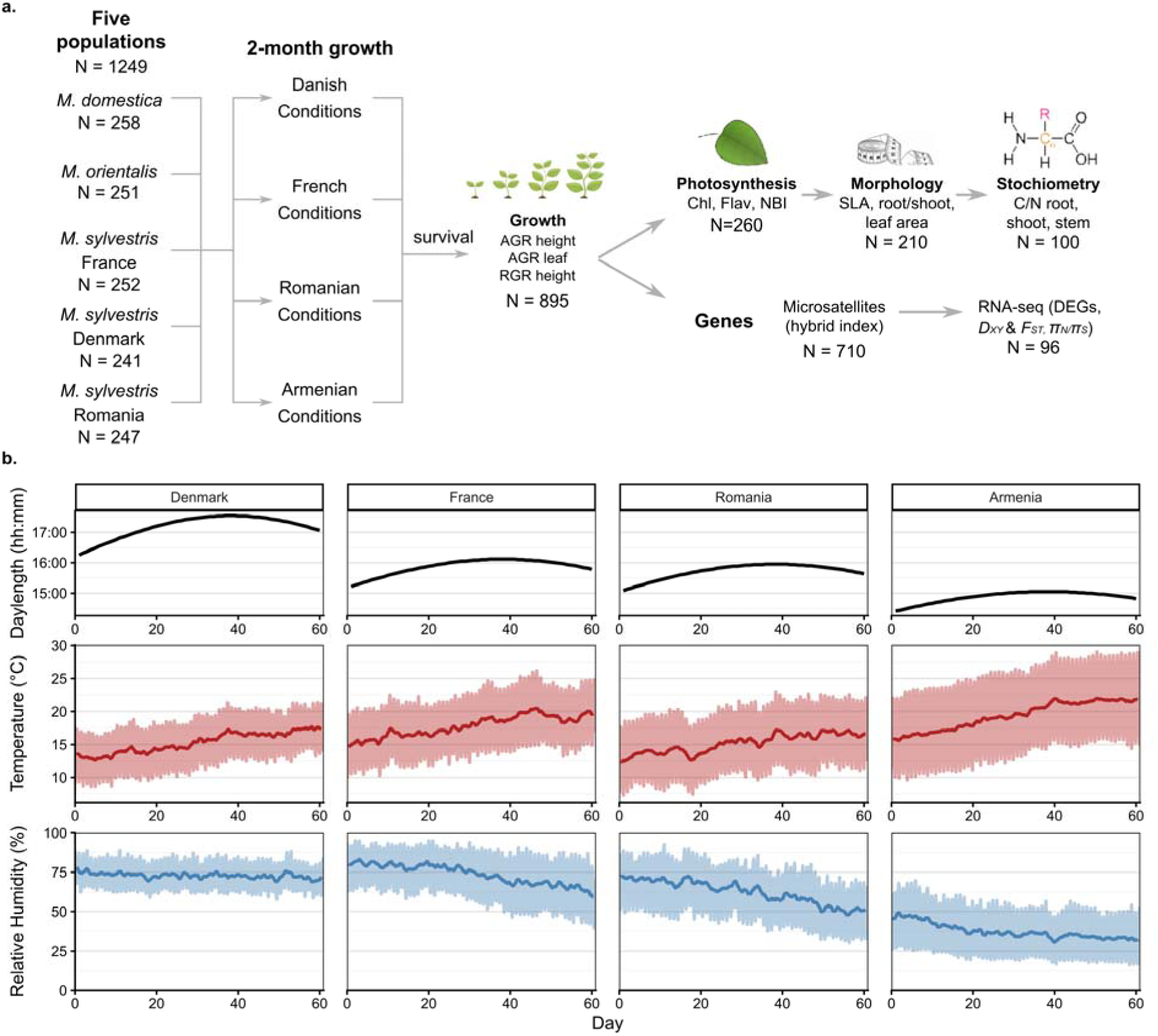
Experimental design of the reciprocal climate-chamber experiment and overview of phenotypic and molecular measurements. (a) Overview of the experimental design and associated phenotypic and molecular data. Seedling survival was monitored over the 2-month experiment. Twelve phenotypic traits were measured across four functional categories: growth (absolute and relative growth rates of height and leaf number), photosynthesis (chlorophyll, flavonoids, nitrogen balance index; N = 260), morphology (specific leaf area, root/shoot ratio, leaf area; destructively sampled subset), and stoichiometry (C/N ratios of roots, shoots, and stems; N = 100). Genetic data included microsatellite genotyping of wild seedlings to estimate the hybrid index (Supplementary fig. S1), whole-genome resequencing of *M. sylvestris* populations to assess genome-environment associations and mutation load, and RNA sequencing (N = 96 after filtering for four low-quality samples; supplementary fig. S1) to characterize differentially expressed genes (DEGs) and gene divergence (*D_XY_*and *F_ST_*). (b) Simulated day length, temperature, and humidity profiles across the ecotron climate chambers. Day length profiles were recreated in ecotron chambers in 2019 (Armenia and Denmark simulated conditions) or in 2021 (France and Romania simulated conditions) based on local day lengths. Mean daily temperature and humidity (dark red and dark blue, respectively) were simulated using 3-hour intervals derived from the average of 30 years of climate records for each region and programmed to reflect realistic diurnal and seasonal variation.

## Results

### Climate treatments shape early survival and phenotypic responses

We established 1,249 seedlings from five apple populations (*M. domestica, M. orientalis, and three M. sylvestris* populations) and grew them under four ecotron climate treatments reproducing contrasting European climatic regimes (France, Denmark, Romania, and Armenia; Fig. 1a,b). Of these, 895 seedlings survived (Fig. 2) and were phenotyped for 12 morphophysiological traits, whereas RNA sequencing was successfully performed on 96 representative seedlings spanning all population × climate treatment combinations (supplementary tables S1-S3).

**Figure 2.**
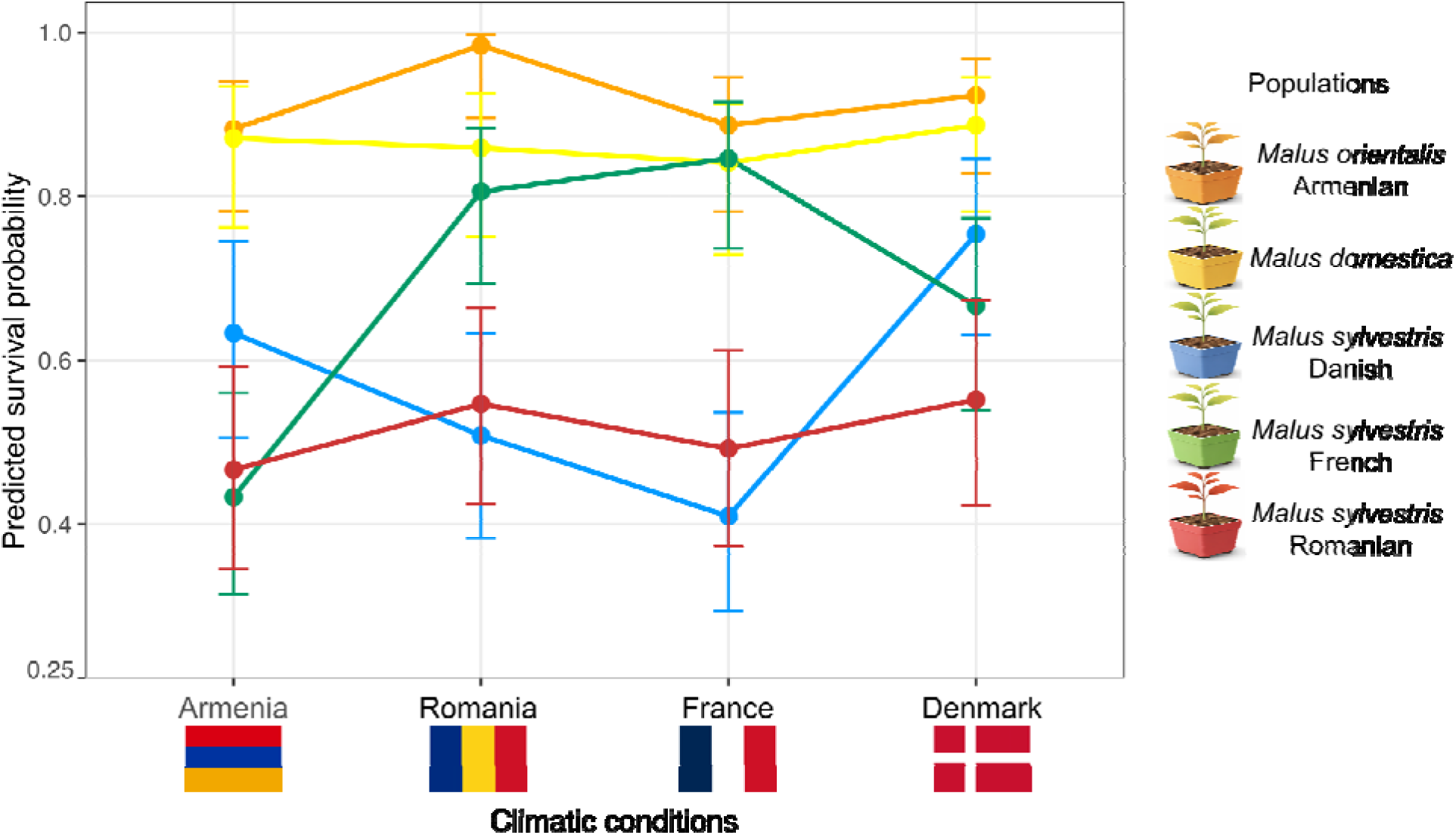
Population origin and climate treatment jointly shape early-stage survival in apple seedlings. Population-specific survival responses across climate treatments. Predicted survival probabilities (±95% CI) at day 61 for five populations (*M. domestica*, *M. orientalis*, and three *M. sylvestris* populations from France, Denmark, and Romania) across four climate treatments. Estimates are derived from the mother-aggregated binomial model (supplementary table S2 and supporting Information Text S1).

Microsatellite markers provided limited resolution for discriminating populations, but enabled robust estimation of crop-to-wild admixture (hybrid index), which was accounted for in downstream phenotypic analyses (supplementary figs. S1-S2). In contrast, RNA-seq-derived SNPs clearly resolved the predefined populations, with pairwise *F_ST_* estimates and neighbor-net analyses supporting strong genetic differentiation among all groups (Fig. 3e; supplementary fig. S1).

**Figure 3.**
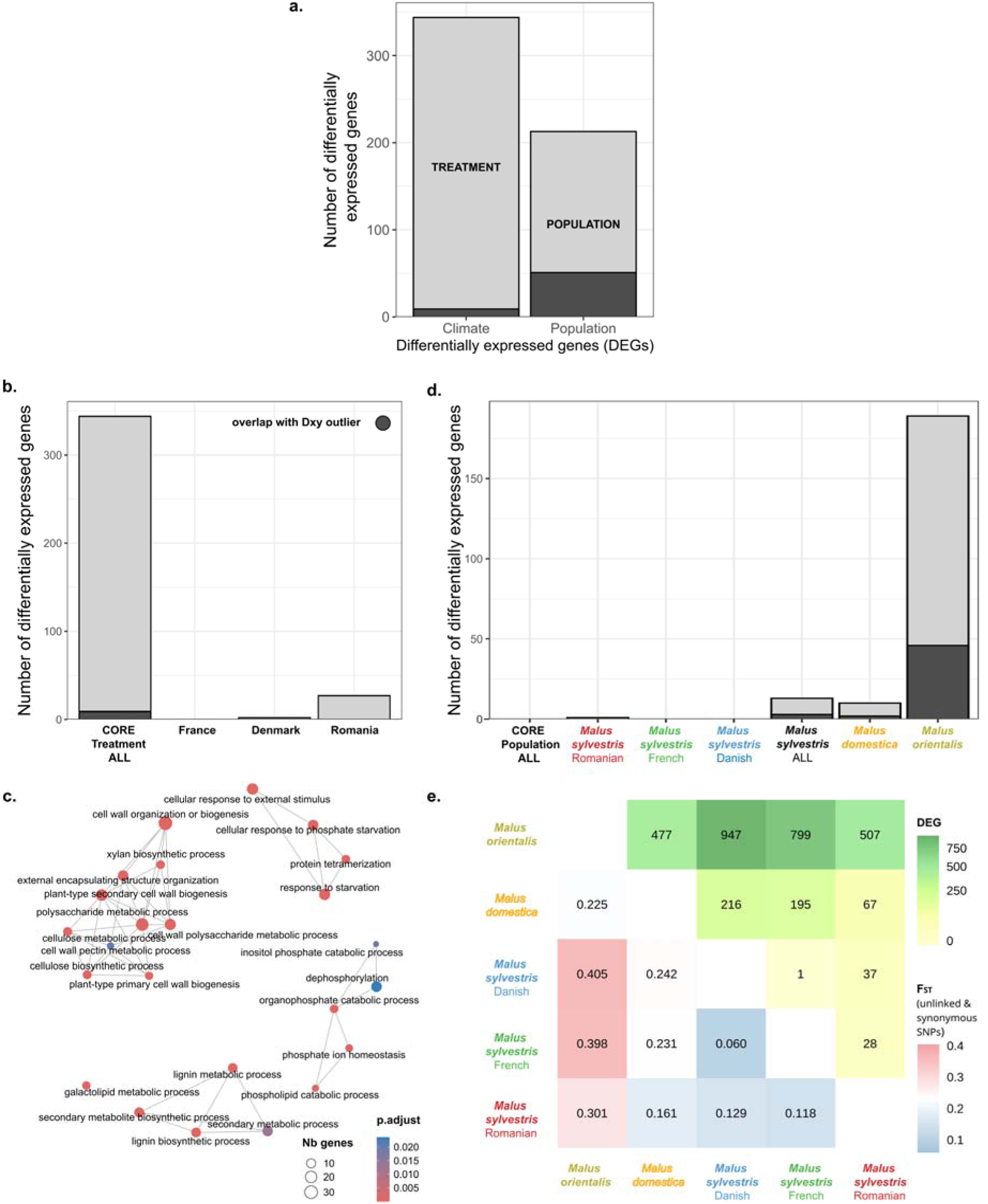
Population and climate treatment effects on gene expression and adaptive variation. (a) Number of differentially expressed genes (DEGs) attributed to the fixed effects of climate treatment or population. (b) Number of DEGs showing a consistent response to climate treatment across all populations (core climate-treatment DEGs), and number of DEGs specific to each climate treatment (France, Denmark, Romania). (c) Gene Ontology (GO) term enrichment map of the 344 core climate-treatment DEGs, showing significant functional enrichment in nutrient response, starvation, cell wall biogenesis, and regulation of seed development. Node size reflects the number of DEGs associated with each GO term; color reflects adjusted p-values. (d) Number of population-specific DEGs per taxon, defined as genes showing consistent expression divergence of one population relative to the others across climate treatments, extracted from population contrasts in the full factorial expression model. Lines connect GO terms with semantic similarity > 0.7, calculated using the Wang method; edge width is proportional to semantic similarity and node size indicates the number of associated DEGs. (e) Heatmap of pairwise population *F_ST_*(bottom triangle, computed using 468,714 synonymous and unlinked SNPs) and number of population-specific DEGs (top triangle) between populations. In (a), (b), and (d), dark grey indicates the proportion of DEGs that overlap with *D_XY_* outliers, highlighting genomic divergence.

Seedling survival was strongly structured by population origin and also varied across climate treatments (Fig. 2; supplementary table S2). Global likelihood-ratio tests revealed a strong population effect (LRT = 165.16, df = 4, *P* < 0.0001), a significant climate-treatment effect (LRT = 10.49, df = 3, *P* = 0.0148), and a significant population × climate-treatment interaction (LRT = 45.59, df = 12, *P* < 0.0001), indicating that populations differed in their survival responses across climate treatments. *M. domestica* and *M. orientalis* maintained consistently high survival across environments, whereas *M. sylvestris* populations showed greater variation. French *M. sylvestris* showed its highest survival under its home-climate treatment, whereas

Romanian *M. sylvestris* consistently exhibited relatively low survival across climate treatments. French *M. sylvestris* survived significantly better under its home-climate treatment, whereas Romanian *M. sylvestris* consistently exhibited the lowest survival across all climate treatments.

AGR height, absolute growth rate of seedling height (cm/day); RGR height, relative growth rate of height, standardized by initial size; AGR leaf, absolute growth rate for number of leaves per day; Chlorophyll, leaf chlorophyll content, as proxy for photosynthetic capacity; Flavonoids, leaf flavonoid content, an indicator of stress response; NBI, nitrogen balance index, calculated as the ratio of chlorophyll to flavonoids; C/N roots, C/N leaves, C/N stem, carbon-to-nitrogen ratio in root, leaf, and stem tissues, respectively; Root/shoot, biomass allocation ratio between root and shoot; leaf area, total surface area of leaves per plant; SLA, specific leaf area (cm²/g), an indicator of leaf morphology and growth strategy.

Climate treatment also emerged as the dominant driver of phenotypic variation (Table 1). Linear mixed models showed significant effects of climate treatment for all twelve morphophysiological traits, whereas population effects were more variable, and significant population × climate treatment interactions were detected for only three traits (relative growth rate in height, absolute growth rate in leaf number, and chlorophyll content), indicating that most phenotypic responses were shared across populations.

**Table 1.**
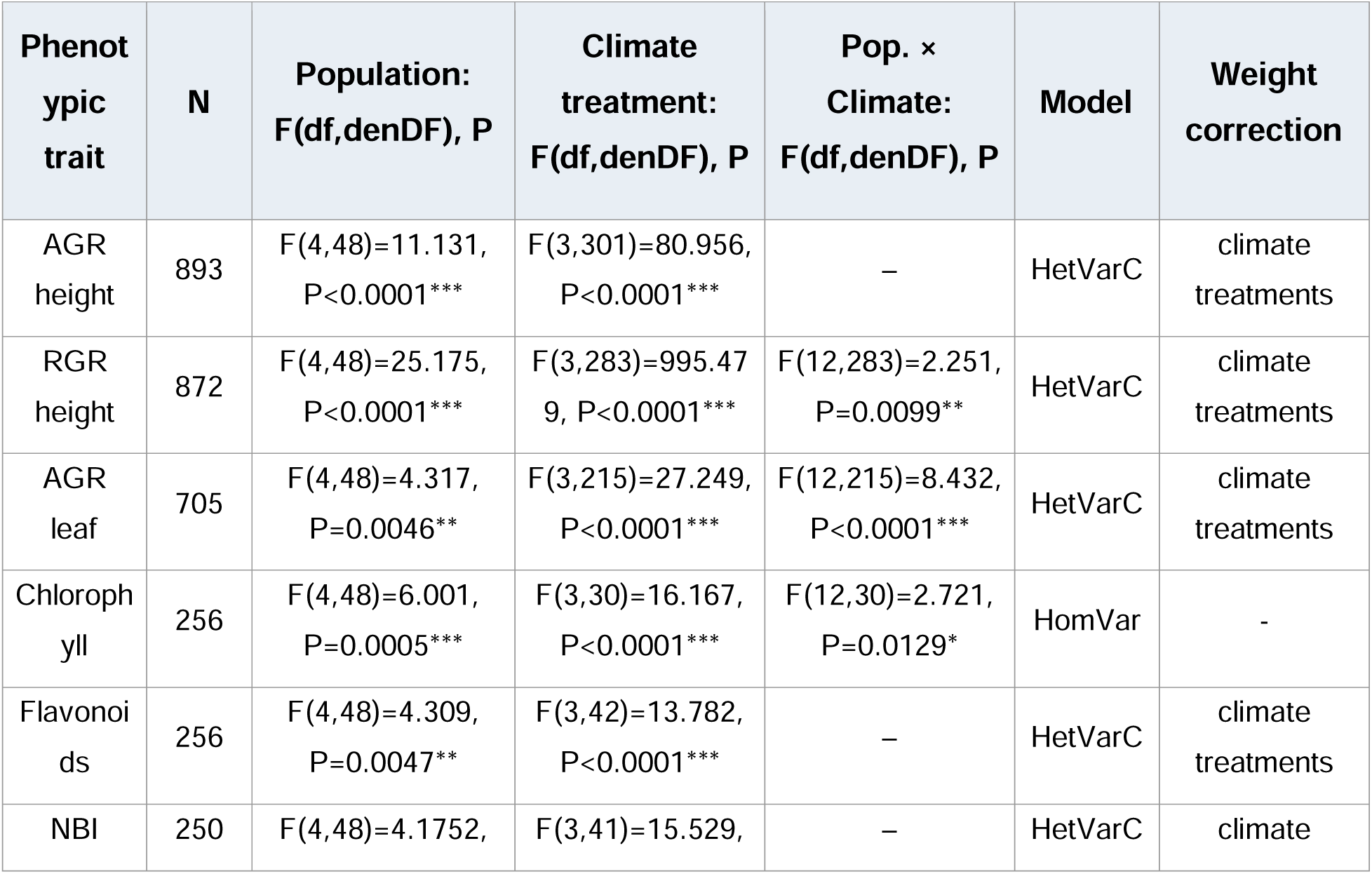
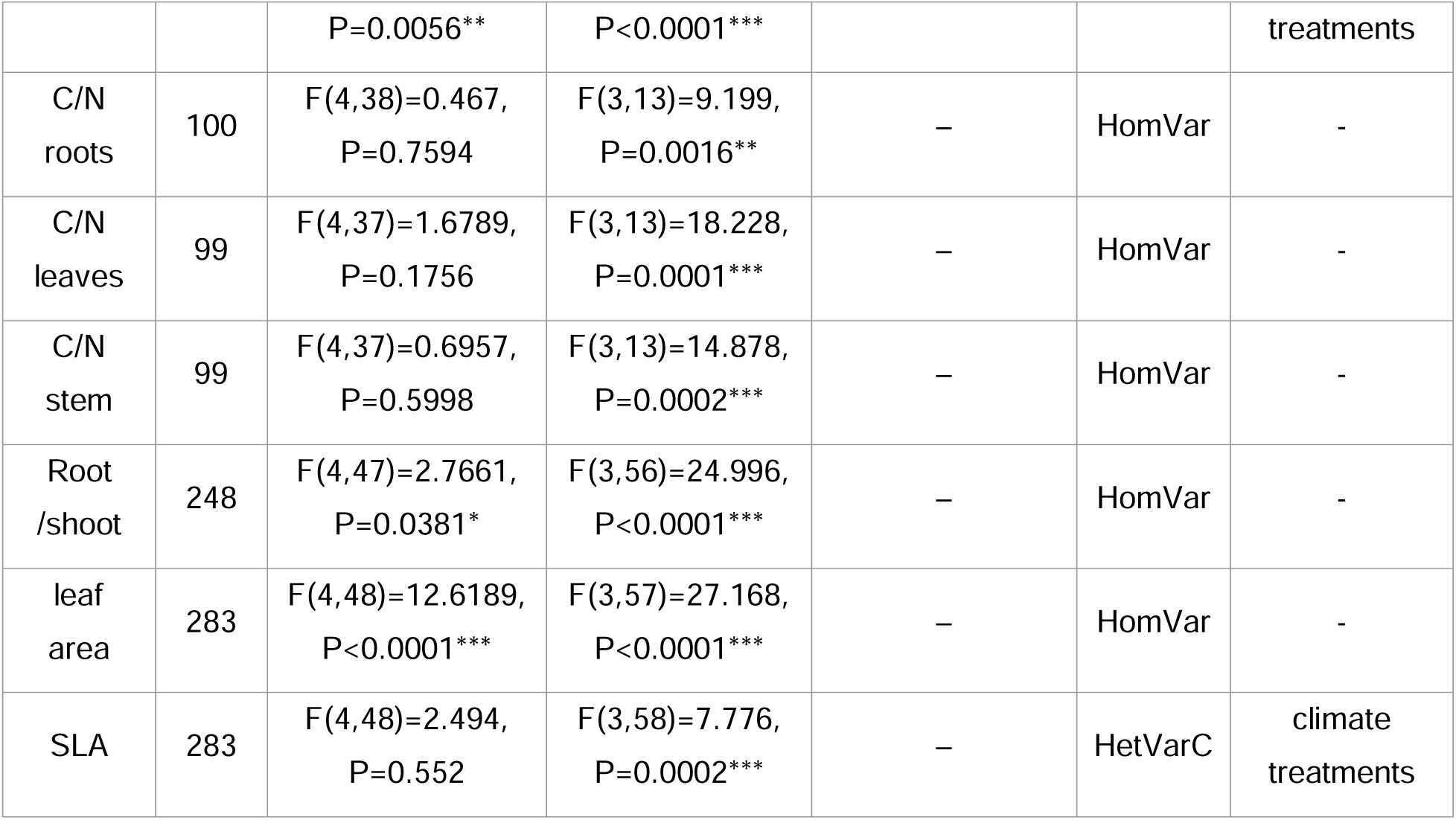
Effects of population, climate treatment, and their interaction on 12 phenotypic traits. . Effects of population, climate treatment, and their interaction on the variation of the 12 phenotypic traits in apple seedlings (linear mixed model). A total of 895 seedlings were grown across four ecotron climate chambers representing climate treatments for Armenia, Denmark, France, and Romania, with experiments conducted in two phases: 2019 (Armenia and Denmark climate treatments) and 2021 (France and Romania climate treatments). Seedlings came from five population groups: cultivated *Malus domestica*, wild *M. orientalis* (Armenia), and *M. sylvestris* from Denmark, France, and Romania. The most appropriate model of residual variance structure is indicated as either HomVar (homogeneous variance) or HetVarC (heterogeneous variance by climate treatments). F-values and associated degrees of freedom (Df, denDF) are reported for each fixed effect (population, climate treatments, and their interaction). Statistically significant results are marked with asterisks (one asterisk: p < 0.05; two asterisks: p < 0.01; three asterisks: p < 0.001). Model-based residual variance corrections are listed in the final column. A dash (–) indicates that the population × climate treatment interaction was non-significant in the initial full model and was therefore removed from the final model.

Climate treatment consistently shaped seedling physiology (supplementary fig. S3). The French climate treatment promoted vigorous shoot growth and larger leaf area, whereas the Armenian climate treatment induced the strongest stress-related responses, including elevated flavonoid accumulation and nitrogen balance index. Population-specific differences were comparatively modest: *M. domestica* displayed larger leaf area, whereas Romanian *M. sylvestris* accumulated higher flavonoid levels. Principal component analysis nevertheless revealed extensive overlap among populations and climate treatments (supplementary fig. S4), indicating that climate primarily induced parallel shifts within a continuous phenotypic space rather than distinct phenotypic clusters.

Random-slope regression mixed models further showed that climate-induced plasticity was widespread but varied among traits and populations (supplementary table S4). Growth traits, chlorophyll content, and root/shoot ratio exhibited significant plasticity in most populations, whereas C/N ratios and specific leaf area showed more population-specific responses. Estimated marginal means further indicated that the Armenian climate treatment consistently elicited stress-related responses, whereas the French climate treatment promoted vegetative growth and biomass accumulation (supplementary tables S4b,c). Importantly, *M. domestica* exhibited substantial phenotypic plasticity across climate treatments, indicating that the magnitude of environmental responsiveness remained comparable to that observed in wild populations. Overall, these results indicate that early climate responses are dominated by widespread climate-induced plasticity shared across populations. We next asked whether this conserved phenotypic response was mirrored by a similarly conserved transcriptional program.

### Climate treatment induces a strong and modular transcriptomic response

Initial transcriptomic analyses across all four climate treatments identified 1,810 core climate-treatment DEGs. However, samples from the Armenian climate treatment were collected at a different time of day than the other three treatments, introducing a potential diurnal confound in gene expression comparisons (Fig. 2; supplementary fig. S5; see Methods). We therefore repeated the analysis on the three comparably sampled treatments (France, Denmark, Romania), recovering 394 DEGs, of which 344 (87%) were already identified in the full analysis (supplementary fig. S6 and S7). This high overlap confirms that the core transcriptional response is robust and that the Armenian exclusion does not qualitatively alter our conclusions. All subsequent analyses are based on these three treatments unless otherwise stated; the Armenian climate treatment is retained for phenotypic and survival analyses, where diurnal sampling is not a concern.

These 344 DEGs were consistently responsive to climate treatment across all populations (Fig. 3a,b; supplementary table S5), highlighting the conserved nature of transcriptional plasticity in response to environmental variation. Fisher’s exact test for overlap with *D_XY_* outliers showed significant depletion (FDR = 2.66 × 10⁻□, odds ratio = 0.27), reinforcing their conserved status. Notably, the Romanian climate treatment induced the most distinct transcriptomic signature, with 27 DEGs unique to this environment (Fig. 3b), compared with 2 and 0 DEGs for the Denmark and France climate treatments, respectively (supplementary table S5). Because RNA-seq was performed exclusively on leaves, these analyses capture only the subset of evolutionary signals associated with genes expressed in this tissue and developmental stage. Accordingly, the absence of overlap between DEGs and genome-wide evolutionary signals should not be interpreted as evidence that a locus is unrelated to climate adaptation.

GO term enrichment analysis revealed that the 344 genes were significantly enriched for functions related to nutrient starvation responses, cell wall biosynthesis, secondary metabolite biosynthesis, and phosphate metabolism (Fig. 3c). Three of these DEGs were also linked to flowering-time regulation (supplementary tables S5 and S6), pointing to environmentally responsive regulation of developmental timing. WGCNA identified 24 co-expression modules, of which two (red and black) were significantly enriched for core climate-treatment DEGs (supplementary fig. S8). These modules showed complementary functional enrichment (supplementary fig. S9): the red module was enriched for cell wall biosynthesis and polysaccharide metabolism, while the black module was enriched for responses to nutrient starvation and phosphate deprivation, suggesting that conserved climate-responsive genes coordinate both structural remodeling and nutrient sensing. Both modules were significantly correlated with multiple phenotypic traits (supplementary fig. S10): the black module showed the strongest associations, with significant negative correlations with growth-related traits (AGR height, RGR, root-to-shoot ratio) and positive correlations with flavonoid accumulation and specific leaf area, while the red module was positively correlated with chlorophyll content and nitrogen balance index. These associations directly link the conserved transcriptional response to measurable physiological and growth variation across climate treatments.

### Limited differential gene expression between populations mostly reflects neutral genomic divergence

In contrast to the strong effects of the climate treatment, half as many genes were differentially expressed among populations (Fig. 3a), and no gene could separate all populations. Nevertheless, some DEGs were found in one population but not in the others (population-specific DEGs). Of these, *M. orientalis* exhibited the largest number of population-specific DEGs (Fig. 3d), consistent with its marked transcriptomic divergence and strong neutral genomic differentiation (Fig. 3e). Importantly, the population-specific DEGs shown in Fig. 3d represent constitutive expression differences among populations across climate treatments and are distinct from the candidate local-adaptation expression patterns presented in Fig. 5b. The latter were identified within *M. sylvestris* by asking whether expression changes were consistently aligned with each population’s home climate treatment (see Methods).

Pairwise DEG counts closely mirrored genome-wide neutral differentiation. Comparisons between *M. orientalis* and *M. sylvestris* populations followed a geographic gradient, with Danish, French, and Romanian *M. sylvestris* populations showing progressively fewer DEGs relative to *M. orientalis* (Fig. 3e). Across all pairwise comparisons, the number of DEGs was strongly correlated with genome-wide *F_ST_* (Spearman’s ρ = 0.95, *P* < 2.2 × 10⁻¹), indicating that population-level expression divergence largely reflects neutral genetic divergence rather than adaptive differentiation.

Only ten population-specific DEGs distinguished *M. domestica* from the wild populations (Fig. 3d), with the fewest differences observed between *M. domestica* and Romanian *M. sylvestris* (Fig. 3e). These included notable candidates such as MD06G1132200, a homolog of the *Arabidopsis thaliana* gene ICE1 (INDUCER OF CBF EXPRESSION 1), and MD15G1374500, a homolog of the *A. thaliana* gene ATHB16 (HOMEOBOX PROTEIN 16) (supplementary tables S5 and S6). Finally, population-specific DEGs significantly overlapped with highly divergent genomic regions (*D_XY_* outliers), particularly in *M. orientalis* (33 of 187 DEGs; FDR = 3 × 10⁻¹¹, odds ratio = 3.89; supplementary table S7), suggesting that although most expression divergence reflects neutral population history, a subset of leaf-expressed genes is associated with genomic regions showing elevated evolutionary differentiation.

### Gene expression divergence and selective constraint in cultivated apple and its wild relatives

Because the core climate-responsive transcriptional program was shared across wild and cultivated apples, we next asked whether domestication reshaped the selective constraints and nucleotide diversity within the coding sequences of the genes involved in this response. Three gene categories have been defined: background genes, core climate-treatment

DEGs (the 344 core climate-responsive DEGs), and dom-wild DEGs (DEGs that distinguish domesticated from wild apples). Across these categories, no significant interaction was detected between population and gene category for log(π*N*/π*S*) (supplementary table S8a), whereas both main effects were highly significant (supplementary table S8a; P < 2.2 × 10⁻¹). Core climate treatment DEGs exhibited the lowest log(π*N*/π*S*) ratio (emmean = −0.4372) (supplementary table S9; Fig. 4a), followed by background genes (−0.2847), indicating stronger purifying selection on climate-responsive genes (supplementary table S9). In contrast, dom-wild DEGs showed the highest ratio (0.0997), consistent with relaxed selective constraint or positive selection. However, variation in π*N*/π*S* alone cannot distinguish among relaxed purifying selection, positive selection, balancing selection, or other selective processes. We therefore interpret these differences conservatively as reflecting shifts in the relative contribution of nonsynonymous versus synonymous diversity, consistent with altered selective pressures.

**Figure 4.**
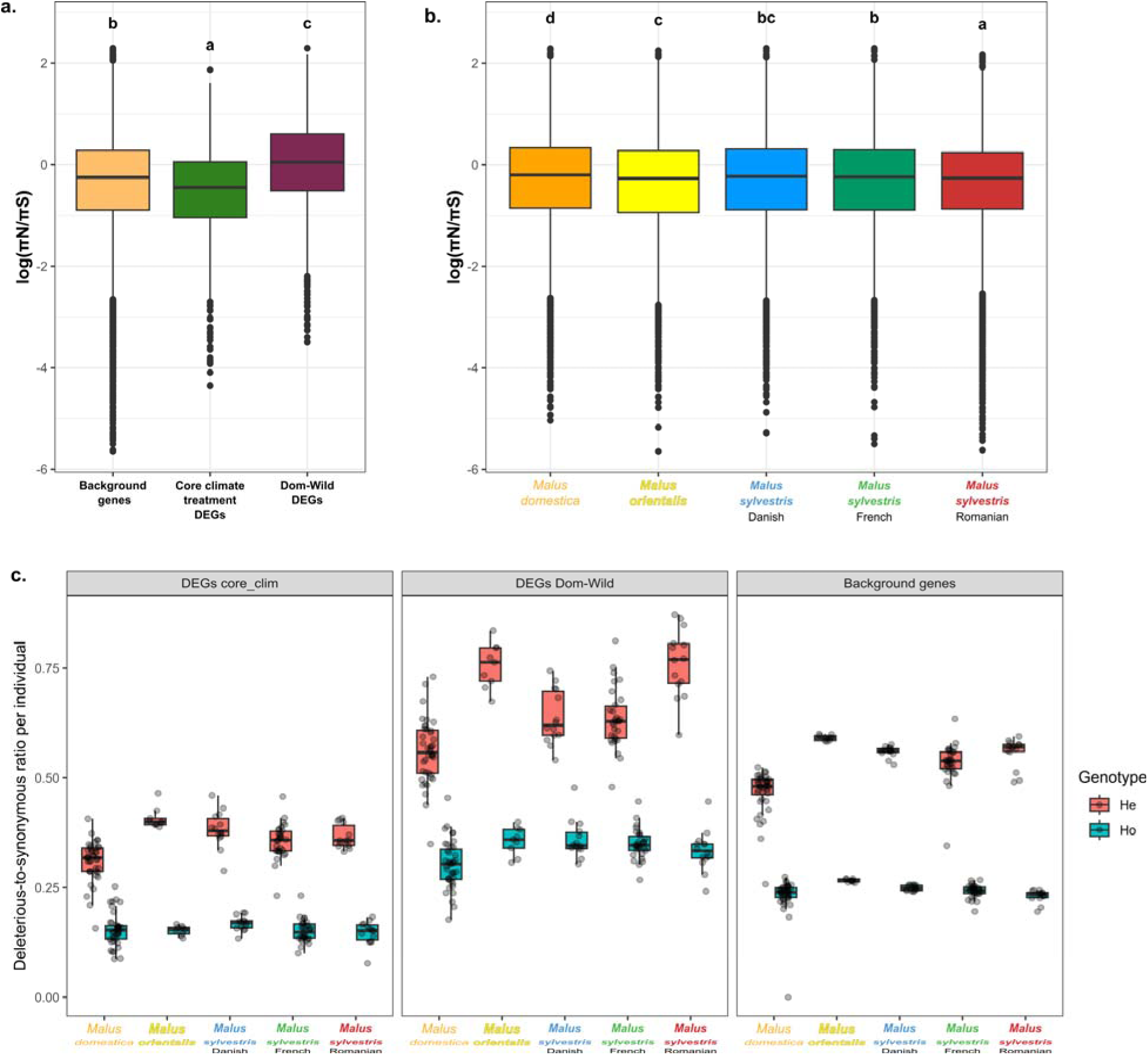
Coding sequence evolution and deleterious mutation load across populations. Patterns of coding sequence evolution and deleterious mutation load across gene categories and apple populations. (a, b) Ratio of nonsynonymous to synonymous polymorphisms (log[π*N*/π*S*]) across (a) gene categories: background genes, core climate-treatment DEGs, and DEGs distinguishing between domesticated and wild apples (dom-wild DEGs) or (b) five apple populations. (c) Deleterious to synonymous mutation ratio per individual for each gene category and population, shown separately for heterozygous (*He*) and homozygous (*Ho*). Genotypes Across populations, *M. domestica* exhibited the highest log(π*N*/π*S*) ratio (supplementary table S10; emmean = −0.168), significantly higher than all wild populations (Fig. 4b; P < 0.0001), consistent with genome-wide relaxation of purifying selection. Conversely, *M. sylvestris*, particularly the Romanian population (emmean = −0.245), exhibited the lowest ratios, suggesting stronger purifying selection.

To explicitly evaluate relative shifts in coding diversity, we modeled the ratio of nucleotide diversity between domesticated and wild apples (π*N_dom_*/π*N_wild_* and π*S_dom_*/π*S_wild_*). These ratios did not differ significantly among the gene categories (π*S* ratio model: P = 0.091; π*N* ratio model: P = 0.056; supplementary tables S8b and S11), indicating that relative shifts in diversity are uniform across coding sequences rather than specific to functional or expression-divergent groups. Consistent with extensive post-domestication introgression from wild relatives rather than a severe domestication bottleneck, the estimated intercepts for both models were significantly positive on the log scale (P < 2 × 10⁻¹□). This demonstrates that *M. domestica* exhibits globally higher nucleotide diversity than wild lineages across all gene groups, confirming that wild-crop gene flow has actively elevated and preserved standing genetic variation in the cultivated lineage.

Patterns of deleterious mutation load paralleled these evolutionary trends. Deleterious burden was quantified as the odds of a mutation being deleterious relative to synonymous (He_del/He_syn and Ho_del/Ho_syn) using a binomial GLMM (Eqn 10). Both Population and GeneType significantly affected relative deleterious burden in both He (*P* = 2.14 × 10^-12^) and Ho (P = 1.37 × 10^-6^) states (supplementary table S13). Core climate treatment DEGs consistently showed the lowest relative deleterious burden, reinforcing their strong evolutionary constraint (Fig. 4c). Across populations, *M. domestica* showed the lowest relative heterozygous (masked) deleterious burden compared to all wild populations. In the homozygous (realized) state, this reduction was significant only for dom-wild DEGs; *M. domestica* did not differ significantly from *M. sylvestris* for background genes or core climate DEGs (supplementary table S13). After normalization by synonymous variation, Romanian *M. sylvestris* was indistinguishable from other *M. sylvestris* populations in both He and Ho burden (all pairwise P > 0.55). *Malus orientalis* and Danish *M. sylvestris* showed comparatively higher homozygous (realized) deleterious burden, consistent with smaller effective population sizes.

Overall, domestication reshaped global patterns of coding sequence diversity, selective constraint, and deleterious variation without altering the conserved climate-responsive transcriptional program. Climate-responsive genes remained under the strongest purifying selection and presented a lower relative deleterious burden than the other gene categories across both wild and cultivated apple lineages.

### Genomic basis of climate adaptation in *M. sylvestris*

To investigate the genomic basis of local climate adaptation in European wild apples, we re-ran our DEG analysis exclusively on the *M. sylvestris* samples and leveraged its extensive geographic sampling to integrate these results with genotype-environment association (GEA) data. This combined genomic and transcriptomic approach allowed us to filter broad evolutionary signals and isolate candidate genes underlying regional climate adaptation.

After filtering correlated environmental variables, LFMM and Samβada identified partially overlapping climate-associated loci, with stronger convergence at the gene level than at the SNP level (supplementary figs. S11-S13), consistent with their different statistical assumptions. The strongest associations involved precipitation (BIO12), followed by temperature-related variables (BIO1, BIO2, BIO3, BIO4, and BIO8; Fig. 5a; supplementary table S14).

**Figure 5.**
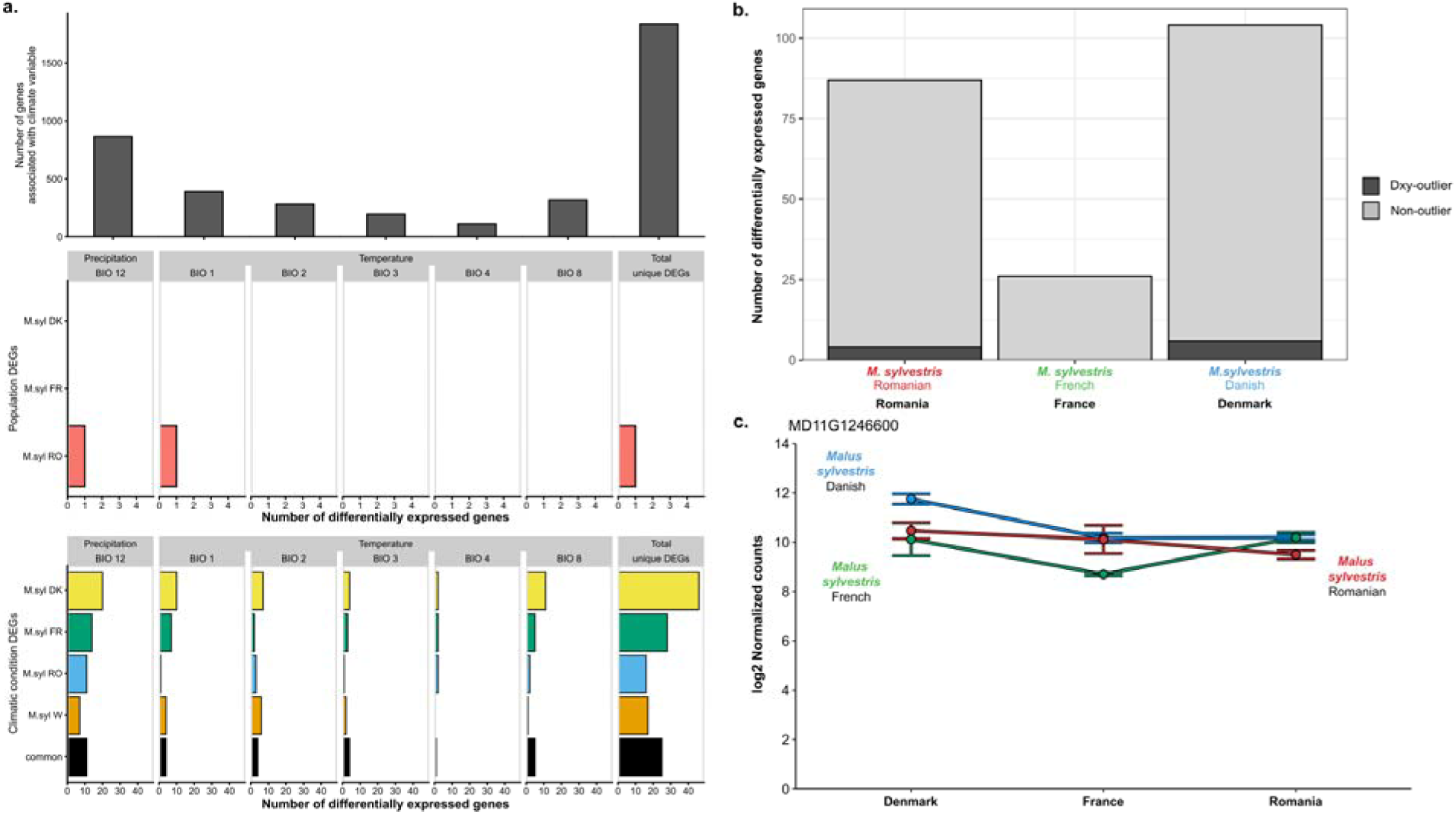
Genomic basis of climate adaptation in *Malus sylvestris*. (a) Intersection between gene×environment associations (GEAs) and differential expression analysis across bioclimatic variables and DEG categories in *M. sylvestris*. Top panel: number of genes significantl associated with each climatic variable (bioclim; p < 0.001) based on both LFMM and Samβada. GEA. The strongest associations were detected for the precipitation-related variable BIO12, followed by temperature-related variables. Middle panel: proportion of population-specific DEGs in *M. sylvestris* that overlap with genes associated with each bioclim variable. Bottom panel: proportion of climate-treatment DEGs categorized as specific to each *M. sylvestris* population, specific to the *M. sylvestris* western population (M. syl W; French and Danish populations), and common to all populations (core climate-treatment DEGs), overlapping with GEA-identified genes for each bioclim variable. (b) Number of candidate local-adaptation DEGs identified within each *M. sylvestris* population, defined as genes showing climate-treatment-aligned expression, i.e., consistently higher or lower expression in the population’s home climate treatment relative to the other climate treatments. Dark grey indicates the proportion of these candidates overlapping *D_XY_* outliers. (c) Log2-normalized expression profile of MD11G1246600 for each population and climate treatment. Each point represents the mean ± standard error; colors represent the respective populations.

Several genes identified by the GEA also overlapped with climate treatment-responsive DEGs, including both core climate-treatment DEGs shared across the three *M. sylvestris* populations and population-specific DEGs (Fig. 5a). This overlap specifically identifies climate-associated loci showing transcriptional responses in leaves at the developmental stage examined here; GEA candidates acting through other tissues or developmental stages would not be detected by our transcriptomic analysis. Across these comparisons, 21 genes overlapped between GEA candidates and climate-treatment DEGs. A permutation analysis showed that this overlap did not exceed random expectation (10,000 permutations; mean expected overlap = 26.9 genes; 95% null interval = 18–37; observed/expected ratio = 0.78; empirical P = 0.90), indicating no significant enrichment of climate-treatment DEGs among GEA candidate genes. To identify candidate genes underlying local adaptation, we further searched for genes showing climate treatment-aligned expression, defined as genes consistently up- or downregulated in their home climate treatment relative to the other climate treatments. This analysis identified 87, 26, and 104 candidate genes in the Danish, French, and Romanian *M. sylvestris* populations, respectively (Fig. 5b; supplementary table S15). Among these, four Danish and six Romanian candidates were also *D_XY_* outliers, indicating concordant transcriptomic and genomic divergence.

Comparison among populations revealed little overlap in candidate genes for local adaptation. No candidates were shared between the French and Romanian populations, whereas six were shared between the French and Danish populations and five between the Danish and Romanian populations. One gene, MD11G1246600, emerged as the strongest candidate (Fig. 5c). It exhibited climate-treatment aligned expression in the Danish population, was identified as a *D_XY_*outlier in both the Denmark–Romania and France–Romania comparisons, and was significantly associated with climatic variables in the GEA. Functional annotation identified MD11G1246600 as a homolog of the *Arabidopsis thaliana* gene *At1g16930*, encoding an F-box/FBD/LRR-repeat protein involved in protein turnover and environmental signaling, supporting a potential role in local climate adaptation.

## Discussion

This study provides the first integrative evidence that both plasticity and genetic adaptation contribute to climate responses in a perennial fruit tree. By combining controlled climate-treatment experiments with transcriptomic and genomic analyses in *M. domestica* and its wild relatives, we dissected the molecular, phenotypic, and evolutionary mechanisms underlying climate responses. Despite the substantial climatic differences among the four ecotron treatments, we observed remarkably conserved transcriptional responses across wild and cultivated apple lineages, whereas evidence of local adaptation was comparatively limited and primarily detected in *M. sylvestris*. Rather than indicating a weak adaptive signal, this contrast suggests that evolutionarily conserved transcriptional plasticity represents the predominant mechanism underlying early climate responses observed in this study. Population-specific regulatory responses and genotype–environment associations nevertheless reveal that local adaptation fine-tunes these responses within particular wild populations. In parallel, domestication was associated with changes in gene expression, reduced deleterious mutation load, and shifts in patterns of molecular diversity, indicating that it reshaped the genomic context in which climate responses evolve rather than fundamentally altering this conserved transcriptional program. Together, these findings provide a refined framework for predicting climate resilience and identifying both conserved and lineage-specific targets for the conservation and breeding of climate-resilient perennial crops.

In perennial species, phenotypic plasticity is increasingly recognized as a key mechanism promoting persistence under environmental uncertainty (Liepe et al. 2022; Kremer et al. 2025; Maury and Plomion 2025). Our results extend this framework to fruit trees, showing that early climate responses are largely driven by widespread yet trait-dependent plasticity. Growth-related traits showed remarkably consistent responses across climates, whereas stress-related physiological traits, such as flavonoid accumulation, responded primarily to climatic harshness. Most wild populations maintained relatively stable phenotypes across environments, consistent with robust plasticity, whereas the Romanian *M. sylvestris* population exhibited reduced stability and poor performance, suggesting demographic constraints or maladaptation. *Malus domestica* also showed broad phenotypic responsiveness across climate treatments, indicating that domestication has not substantially reduced environmental responsiveness in the cultivated lineage. (Cornille et al. 2012; Migicovsky et al. 2021). Collectively, these results identify phenotypic plasticity as the predominant early response to climatic variation in apple seedlings.

At the molecular level, we identified 344 core climate-responsive genes showing remarkably consistent expression shifts across climate treatments and populations. These genes were enriched for stress- and nutrient-related pathways and evolved under strong purifying selection, supporting the existence of an evolutionarily conserved molecular toolkit underlying climate responsiveness, consistent with observations in other tree species (Bräutigam et al. 2013; Maury and Plomion 2025). The fact that this transcriptional program was conserved across both wild and cultivated lineages, despite their contrasting evolutionary histories, suggests that it has been maintained despite the profound demographic and selective changes associated with domestication. The widespread conservation of these transcriptional responses mirrors the broad phenotypic plasticity observed across lineages and indicates that this conserved transcriptional program represents the predominant molecular response to climatic variation in apple seedlings. These findings further support recent syntheses proposing that phenotypic plasticity constitutes a major component of climate resilience in long-lived tree species (Liepe et al. 2022; Maury and Plomion 2025).

Alongside this broadly conserved plastic response, we detected more limited but biologically meaningful signatures of local adaptation. *Malus sylvestris* from France exhibited significantly higher survival under its native-climate treatment and harbored candidate genes with climate-responsive expression, genotype–environment associations, and elevated genomic differentiation, consistent with population-specific adaptation. By contrast, the Romanian *M. sylvestris* population consistently displayed the lowest survival across all climate treatments and reduced phenotypic stability, suggesting maladaptation or demographic constraints rather than local climatic specialization. In comparison, both *M. domestica* and *M. orientalis* maintained high and stable survival across climate treatments, indicating broad environmental tolerance during early establishment. We note, however, that early survival may partly reflect differences in stratification requirements, as stratification conditions were standardized across populations and may have been more favorable for some lineages than others. Nevertheless, the specific survival advantage of the French *M. sylvestris* population under its home climate treatment, together with the convergence of transcriptomic, genomic, and genotype–environment association signals, is consistent with a contribution of local adaptation beyond differences in germination requirements. Together, these results indicate that local adaptation contributes to climate responses in apple but remains comparatively restricted to a subset of *M. sylvestris* populations, where it appears to fine-tune the broadly conserved transcriptional program shared across wild and cultivated apples. Among the candidate loci, MD11G1246600 provides a particularly compelling example, as it combined climate-treatment-aligned expression in the Danish population with elevated genomic differentiation and genotype–environment associations. Functional annotation identified this gene as a homolog of the *A. thaliana* gene

At1g16930, which encodes an F-box/FBD/LRR-repeat protein involved in protein turnover and environmental signaling, providing a plausible functional link to environmental responses. However, because the phenotypic traits measured here represent early-life performance rather than lifetime fitness, these associations remain candidate mechanisms rather than direct evidence of adaptive fitness effects. Long-term reciprocal transplant or common-garden experiments integrating survival, reproductive performance, and targeted physiological or functional assays will be required to determine whether these candidate loci contribute to fitness under contrasting climatic conditions.

Several lines of evidence suggest that the conserved transcriptional plasticity identified here may contribute to climate responsiveness in apple. Climate-responsive genes were enriched for stress- and nutrient-related functions, were evolutionarily conserved, and were under strong purifying selection, indicating that these responses have been maintained over evolutionary time. In addition, a subset of genes showing climate-responsive expression also overlapped with genotype–environment association candidates linked to temperature and precipitation gradients, connecting historical signatures of selection with contemporary transcriptional responses. The relatively limited overlap between GEA candidates and climate-responsive DEGs was not greater than expected under a permutation-based null model, indicating no detectable enrichment of climate-responsive genes among loci associated with long-term climatic variation. This limited convergence may reflect the distinct biological scales captured by these approaches: GEA integrates historical population-level responses to climatic variation, whereas differential expression captures contemporary, leaf-specific transcriptional responses during early development. Differences in statistical power, including the relatively limited sample size of the *M. sylvestris* GEA dataset, may further contribute to the limited overlap. Thus, historical genomic adaptation and contemporary transcriptional plasticity appear to capture largely distinct, although potentially complementary, components of climate response. Similarly, the overlap between population-specific DEGs and loci showing elevated *D_XY_* may indicate convergence between expression and sequence differentiation, but does not by itself demonstrate adaptive regulatory divergence. Such associations could reflect direct cis-regulatory differentiation, linkage or hitchhiking with nearby selected variants, or other effects of population history, and distinguishing among these alternatives would require fine-mapping and functional validation. Moreover, our transcriptomic analyses were restricted to leaves and therefore capture only climate-responsive genes expressed in this tissue and developmental stage. Climate adaptation in perennial trees may also involve organ- and stage-specific processes, including root responses, bud dormancy, flowering, seed physiology, and reproductive development, which were not captured by our experimental design. Consequently, the absence of overlap between DEGs and genome-wide evolutionary candidates should not be interpreted as evidence for the absence of an adaptive role. Together, these observations are consistent with the hypothesis that part of this conserved response has been shaped by selection and may contribute to environmental performance.

Common-garden experiments frequently reveal genotype × environment interactions and genetically variable reaction norms for ecologically important traits (Rehfeldt et al. 2002; Chambel et al. 2005; Marchi et al. 2022; Leites and Benito Garzón 2023). However, demonstrating adaptive plasticity requires evidence that plastic responses increase fitness across environments, which remains particularly challenging in long-lived perennial species because lifetime reproductive success is rarely measurable (McGraw and Caswell 1996). Although early vegetative traits, such as growth and flavonoid accumulation, serve as informative proxies for physiological performance under climatic stress, they cannot fully capture lifetime fitness. Long-term common-garden or reciprocal transplant experiments integrating reproductive output, survival, carbon assimilation, and repeated climatic stress will therefore be essential to determine the adaptive value of these plastic responses (Nicotra et al. 2010;

Hartmann et al. 2020). Our results thus provide strong evidence for widespread and evolutionarily conserved transcriptional plasticity, while highlighting that its adaptive significance remains an important question for future studies.

Domestication also influenced the genomic context in which climate responses evolve. Despite exhibiting broad and stable phenotypic performance across climate treatments, *M. domestica* retained the conserved climate-responsive transcriptional program shared with its wild relatives, indicating that domestication did not fundamentally alter the core transcriptional response. Instead, domestication was primarily associated with changes in coding sequences patterns of molecular diversity and gene expression. Although the core climate-responsive genes remained under strong purifying selection and carried consistently low relative deleterious burden, the relative shifts in nucleotide diversity during domestication (π*N_dom_*/π*N_wild_* and π*S_dom_*/π*S_wild_*) did not differ among functional gene categories, indicating that diversity patterns were driven more strongly by population history than by gene function. Consistent with extensive wild-crop introgression rather than a severe domestication bottleneck, *M. domestica* maintained distinct genome-wide diversity profiles relative to wild populations. However, methodological and sampling effects may also contribute to the observed differences in relative deleterious burden. Deleterious variants were identified using SIFT4G predictions, which provide a consistent annotation framework across *Malus* populations, but prediction accuracy may vary with annotation quality and evolutionary conservation across genes. Moreover, although our gene-set subsampling procedure controlled for differences in gene-set size, differences in population sampling could still influence estimates of deleterious burden. Demographic history and wild-to-crop introgression should therefore be regarded as plausible, rather than exclusive, explanations for the comparatively low relative deleterious burden observed in *M. domestica*.

Nevertheless, cultivated apples exhibited the highest overall log(π*N*/π*S*) ratio, indicating a greater relative contribution of nonsynonymous versus synonymous diversity. Such a pattern is consistent with relaxed purifying selection associated with agricultural management and clonal propagation, which buffer against natural selection (Gaut et al. 2018), but variation in π*N*/π*S* alone cannot distinguish this process from positive selection, balancing selection, or other changes in selective pressures. In addition, genes differentially expressed between cultivated and wild apples (dom-wild DEGs) displayed the highest log(π*N*/π*S*) ratios among all gene categories, indicating that transcriptional divergence associated with domestication evolved under a distinct selective regime. Together, these results indicate that domestication reshaped patterns of molecular diversity and deleterious mutation load, thereby modifying the genomic context in which climate responses evolve rather than fundamentally altering the conserved transcriptional response shared between wild and cultivated apples.

Our results are consistent with the hypothesis that conserved transcriptional plasticity may facilitate subsequent genetic adaptation, as proposed by the Baldwin effect (Crispo 2008; Kremer et al. 2014). Similar interactions between plasticity and genetic adaptation have been reported in long-lived tree species, where multisite common-garden experiments indicate that phenotypic plasticity and genetic differentiation jointly contribute to climate resilience (Ramírez-Valiente et al. 2010; Maes et al. 2019; Alexandre et al. 2020; Caignard et al. 2024). Although our results identify candidate genes that combine plastic expression and signatures of selection, demonstrating that plasticity directly facilitates evolutionary change will require long-term experiments that link plastic responses to fitness and genetic change across generations.

Despite the depth of our dataset, our controlled experiments required compromises, including trade-offs among sampling depth, replication across mother lines, and environmental realism. One limitation of the experimental design is that the four climate treatments were conducted over two experimental years due to logistical constraints with the ecotron facilities.

Experimental year was explicitly accounted for in all statistical analyses, and transcriptomic analyses were repeated after excluding the Armenian treatment, which differed in sampling timing. Although experimental year, sequencing batch, and climate treatment cannot be completely disentangled, the strong concordance between analyses performed with and without the Armenian treatment supports the robustness of the main conserved transcriptional response. Population-specific contrasts, including comparisons between the French and Romanian populations, should nevertheless be interpreted with caution because residual batch × treatment effects cannot be completely ruled out. In addition, climate simulations represented the same calendar window across sites rather than site-specific phenological growing-season onset. Consequently, the experiment captures responses to contrasting climates during a standardized early-season period but does not reproduce geographic variation in the timing or duration of the natural growing season. Further validation in long-term field trials and across developmental stages will nevertheless be valuable (Cornille and Alix 2024; Criado et al. 2026).

The within-group sample size (n = 5 seedlings per population × climate treatment combination) reduces statistical power specifically for detecting population-level differential expression and rare or small-effect regulatory differences between populations. Importantly, even with this limited per-group sample size, our design detected 394 climate treatment DEGs and significant GEA signals in *M. sylvestris*, suggesting that our data were sufficient to capture moderate-to-large effects. Population-specific results should therefore be interpreted with caution, particularly given the limited number of DEGs distinguishing *M. domestica* from wild populations (n = 10), and replication with larger RNA-seq sample sizes would strengthen these conclusions. Finally, our study does not capture epigenetic mechanisms, which may also mediate environmentally induced and potentially heritable variation in long-lived plants (Niederhuth and Schmitz 2014; Quadrana and Colot 2016; Baduel et al. 2025). Integrating epigenomic data will be an important step toward disentangling genetic, transcriptional, and epigenetic contributions to climate adaptation.

Our study demonstrates that early climate responses in apple are dominated by evolutionarily conserved transcriptional plasticity, whereas local adaptation contributes more modestly and primarily within wild populations. Domestication did not fundamentally alter this conserved response but reshaped the genomic context in which it evolves. More broadly, these findings illustrate how integrating phenotypic, transcriptomic, and genomic approaches can disentangle conserved and lineage-specific mechanisms of climate resilience in perennial crops, providing a conceptual framework for predicting responses to future climate change and guiding conservation and breeding strategies.

From an applied perspective, these complementary signals provide distinct entry points for climate-resilience strategies. Core climate-responsive genes could be prioritized for functional validation as candidate expression-based biomarkers of broad climatic plasticity and subsequently tested across germplasm collections and developmental stages. Conversely, climate-associated loci and population-specific adaptive candidates in wild *M. sylvestris* may help identify populations or genotypes carrying potentially valuable adaptive variation for conservation and pre-breeding. Integrating these candidates with phenotypic performance under future climate scenarios could ultimately enable more targeted screening of wild and cultivated germplasm. However, the candidates identified here should not yet be considered predictive markers of climate resilience, as their causal effects on fitness and stress tolerance remain to be experimentally validated.

## Methods

### Experimental design and climate treatments

Seeds from five apple populations were used: *M. sylvestris* from Romania, France, and Denmark (Cornille et al. 2015); *M. orientalis* from Armenia, known for its stress tolerance (Volk et al. 2015); and cultivated apple (*M. domestica*) from controlled crosses between cultivars representing a large diversity from traditional or recent compartments maintained by the Biological Resource Center “RosePom - Pome Fruits and Roses” (supplementary table S1; Fig. 1a, supplementary fig. S1). For each population, 9-12 seeds per mother tree were stratified for three months at 4°C in moist sand (50:50 v:v) and later sown into climate-controlled ecotron chambers (Roy et al. 2021).

To expose the seeds to the historical climate treatments of the four native sites, they were placed in ECOTRON climate chambers programmed to replicate selected temperature and humidity profiles. These specific temperature and humidity setpoints (targets) were calculated by averaging historical data from 1985 to 2014 over 3-hour intervals for the same seasonal window (May 15–July 15), resulting in a total of 488 climate targets per native site. The historical temperature and humidity datasets were retrieved from the Climate Data Interface (CDI). Daylight periods were adjusted daily for each chamber to replicate natural daylight variation from 15th May to 15th July at each native site. To this end, daylight time for each native site was extracted from the ptaff database.

The duration of the experiment, 2 months, was selected based on previous results indicating that the growth rate plateaued after a month and a half (Chen et al. 2023). During this period, trays were rotated daily, and seedlings were irrigated once a week. Due to space and time limitations, the experiment was conducted in two batches (years), with the Armenian and Danish treatments conducted in 2019 and the French and Romanian treatments in 2021 (Fig. 1b, Supporting Text S1).

In total, 1,249 seeds were stratified and sown, and monitored for 2 months, across the four ecotron treatments, resulting in approximately 60-70 seeds per population per climate treatment (supplementary tables S1-S2). Of these, 895 seeds survived after germination/establishment and were used for downstream analyses.

### Measures of survival, growth, and physiological traits

Throughout the experiment, growth traits (number of leaves, plant height) were measured individually every 2 to 3 days for all seeds. Those measurements were used to calculate individual absolute (AGR) and relative (RGR) growth rates for height and leaf number using previously reported formulas (Briggs et al. 1920; Radford 1967).

Photosynthetic traits chlorophyll (Chl), flavonoids (Flav), and the nitrogen balance index (NBI) were measured on the final day of the experiment (day 61) using a DualexⓇ instrument (Force-A, France) on 13 seedlings per population per climate treatment (*N* = 260 in total, Fig. 1a). Finally, sampling of 5 to 19 seedlings per population was used to determine leaf area, root/shoot ratio, specific leaf area (SLA) and carbon and nitrogen content for leaf, stem and root. Carbon and nitrogen content were determined by dry combustion using a CN auto-analyzer (Flash EA 1112 series, Therm Finnigan).

At the end of the experiment, individual survival was recorded (supplementary table S2, Supporting Text S1). Survival was defined as successful emergence and growth in pots, integrating both germination and early seedling establishment, which were not recorded separately. Genomic DNA from surviving *M. orientalis* and *M. sylvestris* seedlings was genotyped using 12 simple sequence repeat (SSR) markers, and a hybrid index (*Pdom*) was computed with the introgress R package (Gompert and Buerkle 2010) (supplementary table S1, Supporting Text S1).

### Survival, growth, and physiological trait analysis

In total, for surviving seedlings, 12 traits were analyzed across four categories: growth (AGR height, AGR leaf, RGR height), photosynthetic (Chl, Flav, NBI), stoichiometric (C/N of stem, leaf, and roots), and morphological (specific leaf area (SLA), root/shoot, leaf area; Table S1) traits. Subsampling and trait overlap ensured consistency across the analyses (supplementary tables S1 and S3).

Seedling survival on the last day of the experiment was analyzed as a binary trait using binomial models including population, climate treatment, and their interaction. To account for non-independence among seedlings from the same mother, survival counts were aggregated by mother identity and analyzed using a mother-aggregated binomial GLM. Global effects of population, climate treatment, and their interaction were evaluated using likelihood-ratio tests based on hierarchical comparisons of nested models while preserving model marginality (supplementary table S2; Supporting Text S1).

Batch (year) effects were tested using a linear mixed model (lme4 (Bates et al. 2015)) (Eqn 1; Supporting Text S1). Growth, photosynthetic, stoichiometric, and morphological trait values were corrected using best linear unbiased predictions (BLUPs) from random-effects models (Eqns 2, 3; supplementary tables S16 and S17; Supporting Text S1). Corrected traits were used in a principal component analysis (PCA) with the FactoMineR package (Lê et al. 2008). Additionally, a linear mixed-effects model was fitted for each individual trait using lme4. Each model included population, climate treatment, and their interaction as fixed effects, and hybrid index and mother identity as random effects. For each trait, the population × climate treatment interaction was first tested in the full model. When the interaction was not significant, it was removed, and the model was refitted with population and climate treatment as the only main effects. For each trait, the population × climate treatment interaction was first tested in the full model. When the interaction was not significant, it was removed, and the model was refitted with population and climate treatment as main effects only. Accordingly, population × climate treatment interaction terms are reported in Table 1 only when retained in the final model.

Heteroscedasticity was assessed by comparing homoscedastic models with models that allow unequal residual variance across years (2019 vs. 2021) using analysis of variance (ANOVA). Final model selection was based on the lowest Akaike Information Criterion (AIC) (see Supplementary Information 1 for further details).

### Local adaptation and plasticity

Local adaptation was assessed using linear mixed-effect models with population, climate treatment, and their interaction as fixed effects, and batch (year) and mother tree as random effects (Eqn 4; Supporting Text S1). Model selection was based on the Akaike information criterion (AIC).

Climate treatment-induced phenotypic plasticity within populations was quantified using random-slope regression mixed models (RRMMs, Eqn 5, (Schielzeth and Nakagawa 2013; Gómez et al. 2025)), with climate treatment treated as a fixed effect and mother as a random effect.

### RNA extraction and gene expression analysis

Leaves from 100 seedlings (20 per population) were sampled on the last day of the experiment (supplementary table S3, Fig. 1a). Leaves were chosen for transcriptomic analyses as they integrate climatic signals related to photosynthesis, water balance, and metabolic regulation. To standardize collected tissue, sampled seedlings were at the 7-leaf stage, and for each, an approximately 2 cm square of the lamina was collected. For each seedling, total RNA was extracted using NucleoSpin (Macherey-Nagel) with the addition of PVP40 and CTAB, and its quality was assessed using a NanoDrop and a Bioanalyzer. RNA-seq libraries were prepared by Novogene with mRNA enrichment and the Novogene NGS Stranded RNA Library Prep Set (PT044 internal kit) and sequenced on a NovaSeq 6000 instrument as paired-end 150-bp reads.

The quality of raw reads was assessed with FastQC (Andrews 2010), and then reads with a mean Phred score lower than 15 were filtered using fastp, with trimming of the first 10 bases due to lower overall quality; SortMeRNA was used to remove remaining ribosomal RNA with default parameters (Kopylova et al. 2012; Chen et al. 2018). After filtering, 96 samples passed QC. Filtered reads were mapped to the GDDH13 reference transcriptome (Daccord et al. 2017) using STAR (Dobin et al. 2013) and quantified with Salmon (Patro et al. 2017).

Differentially expressed genes (DEGs) were identified using the DiCoExpress pipeline (Lambert et al. 2020). Because RNA-seq was performed exclusively on leaf tissue, all DEG-based analyses and subsequent comparisons with genome-wide evolutionary signals are restricted to genes expressed in leaves at the developmental stage sampled here. Differential gene expression analyses were performed twice: first, including all four climate treatments; and second, after excluding the Armenian treatment, which was sampled at a different developmental stage due to experimental constraints. This second analysis removed the only treatment affected by differences in sampling time while retaining the three climate treatments sampled under a common experimental schedule. Throughout the study, the experimental year (2019 vs. 2021) was included as a fixed effect in all statistical models to account for potential batch effects. The DiCoExpress pipeline was run using a false discovery rate (FDR) threshold of 0.05, trimmed mean of M-values (TMM) normalization (Robinson and Oshlack 2010), and a filtering threshold of counts per million (CPM) > 5. Only genes showing an absolute log2 fold change > 1 (equivalent to a fold change > 2) were considered differentially expressed. Generalized linear models (GLMs) were then used to test the effects of population, climate treatment, and their interaction on gene expression (Supporting Text S1).

To examine gene expression variation, normalized gene counts were modeled as a function of population, climate treatment, and their interaction using a general linear model:

GE_ijk_ ∼ μ+Population_i_+ Climate treatments_j_+(Population* climate treatments*)_ij_+*ε*ijk*

Where GE_ijk_ is the normalized expression level of a given gene for a seedling from population i under climate treatment j, μ is the overall mean, and εijk is the residual. The interaction term captures potential genotype-by-climate (G × E) interactions.

Population-associated expression differences were evaluated from the full factorial model described above. Contrasts derived from this model were then used to identify genes showing consistent population-associated expression differences across climate treatments. Genes significant in at least three of the four climate-treatment-specific population contrasts and showing a consistent direction of expression were retained as population DEGs.

Climate treatment DEGs were defined as genes showing expression variation within a given population across treatments. Climate treatment DEGs were subdivided into core DEGs (shared across all populations), species-specific DEGs (specific to *M. domestica*, *M. orientalis*, or *M. sylvestris*), and population-specific DEGs within *M. sylvestris*.

Finally, the sets of DEGs were compared to previously identified genes under positive or balancing selection in wild and cultivated *Malus* populations (Chen et al. 2026), to evaluate whether expression patterns were associated with genomic signatures of selection. Full expression profiles for DEG categories, along with their *D_XY_* outlier status, are detailed in supplementary table S5.

We performed an additional differential gene expression analysis restricted to *M. sylvestris* RNA-seq data, in French, Romanian, and Danish climate treatments, and redefined core climate treatment genes as those consistently differentially expressed across populations in response to climate treatment, reflecting conserved molecular plasticity. Population-specific DEGs were defined as genes differentially expressed across all climate treatments among populations.

### Gene ontology (GO) term enrichment and co-expression network analysis

GO term enrichment analysis was performed using the R package *clusterProfiler* (v4.6.0) (Xu et al. 2024) with a custom OrgDb from the GDDH13 reference genome. A Benjamini-Hochberg false discovery rate (FDR) correction (Benjamini and Hochberg 1995) was applied. Weighted Gene Co-expression Network Analysis (WGCNA) (Langfelder and Horvath 2008) was used to identify gene modules correlated with phenotypic traits and to intersect them with lists of DEGs. Enrichment analysis was performed on modules containing more than 50 DEGs.

### Genetic divergence estimated using RNA sequencing data

Single-nucleotide polymorphisms (SNPs) were called from RNA-seq reads using GATK HaplotypeCaller (Auwera and O’Connor 2020) and annotated with SnpEff and SnpSift (Cingolani et al. 2012). In total, 1 million exon-wide SNPs were identified from the RNA-seq data (Supporting Text S1, supplementary fig. S14), including 468,714 synonymous SNPs.

To quantify neutral population genomic differentiation, pairwise genetic differentiation (*F_ST_*) among populations was estimated using Pixy (Korunes and Samuk 2021). Analyses were restricted to 468,714 synonymous and unlinked SNPs and invariant sites to minimize the influence of selection and linkage and to avoid measurement biases, following the default Pixy behavior, thereby providing robust estimates of background genomic differentiation among populations.

To investigate whether differential gene expression was associated with elevated genetic differentiation, gene-level absolute divergence (*D_XY_*) was estimated to identify highly divergent genes (supplementary fig. S14). The average number of nucleotide differences per site (*D_XY_*) was first calculated per exon using Pixy(Korunes and Samuk 2021). Gene-level *D_XY_*values were obtained by aggregating the Pixy metrics count_diff and count_comparison across all exons within a gene. Absolute divergence (*D_XY_*) was preferred over relative metrics such as *F_ST_* because it is sensitive to within-population genetic diversity and can be biased by local variation in recombination rates or linked selection (Charlesworth 1998). Gene-level *D_XY_* values were standardized using Z-scores, and genes with Z-scores greater than 2 were classified as highly divergent outliers (Supporting Text S1). The overlap between these genetically divergent genes and DEGs was then assessed to evaluate the contribution of genetic divergence to population-level differences in gene expression.

To evaluate whether specific categories of DEGs were significantly enriched or depleted among *D_XY_* outliers, we performed overrepresentation and underrepresentation analyses using Fisher’s exact tests. For each DEG category, a 2×2 contingency table was constructed based on the presence or absence of genes within that category relative to their status as *D_XY_* outliers. We performed separate one-tailed tests to identify both enrichment (alternative = ‘greater’) and depletion (alternative = ‘less’) of outliers within each category. To account for multiple hypothesis testing across all categories and directions, p-values were adjusted using the Benjamini-Hochberg False Discovery Rate (FDR) method.

### Population genetic differentiation: microsatellite markers and RNA-seq-derived SNPs

Because the populations used in the experiment were defined a priori based on geographic origin and taxonomic assignment (Cornille et al. 2015), we assessed whether these groups were supported by genetic differentiation. The 12 microsatellite (SSR) markers provided limited resolution to discriminate populations, with substantial overlap among groups, as expected given the small number of loci (supplementary fig. S1b,c). We therefore did not use these markers to infer population structure *de novo*. Instead, they were used to estimate individual hybrid ancestry with cultivated apple through the hybrid index (*Pdom*) (Gompert and Buerkle 2010), which was included as a random effect in downstream phenotypic models to control for crop-to-wild admixture (Supporting Text S1). To independently validate genetic differentiation among the predefined groups, we used SNPs called from the RNA-seq data (see section ‘Genetic divergence estimated using RNA sequencing data’ above) and constructed a neighbor-net phylogenetic network using poppr (Kamvar et al. 2014) and phangorn (Schliep 2011) (supplementary fig. S2a). The RNA-seq subset was not restricted to non-admixed individuals; naturally occurring crop-to-wild introgression was retained because it represents standing genetic variation within contemporary wild populations. This differs from the independent WGS dataset used for GEA analyses, in which individuals with *M. sylvestris* ancestry below 0.9 were excluded to ensure association signals reflected wild-population variation (Supporting Text S2). This higher-resolution marker set, comprising 468,714 synonymous and unlinked SNPs, revealed clear separation among all population pairs, providing quantitative support for the biological relevance of the a priori population grouping used throughout the analyses.

### Genetic diversity, gene expression, and selection efficiency

Gene expression variation, molecular diversity, and deleterious mutation load were compared between the cultivated apple (*M. domestica*) and its wild relatives (*M. sylvestris* and *M. orientalis*). Analyses focused on three gene categories: (i) dom-wild DEGs, (ii) core climate treatment-responsive DEGs, and (iii) background expressed genes (Sauvage et al. 2017).

To assess the impact of domestication on molecular diversity, the 1,086,764 SNPs called from RNA-seq data (see above “Genetic divergence estimated using RNA sequencing data” and Supporting Text S1) were used to estimate nucleotide diversity at nonsynonymous (π*N*) and synonymous (π*S*) sites with Pixy (Korunes and Samuk 2021). We modeled the log-ratio of diversity between cultivated and wild apples as a function of gene category for both π*N* and π*S* (Supporting Text S1, Eqns 7-8), thereby distinguishing genome-wide effects of domestication from gene-category-specific deviations. Differences in purifying selection efficiency were further evaluated by modeling π*N/*π*S* across populations and gene categories, including their interaction (Supporting Text S1, Eqn 9).

Derived deleterious alleles and synonymous variants were taken from the whole-genome SNPs dataset of Chen et al. (Chen et al. 2026), where they had been annotated using SIFT 4G (Vaser et al. 2016). Only individuals from populations included in the present study were kept. For each individual deleterious mutation counts were evaluated relative to synonymous counts within each gene category and analyzed using binomial generalized linear mixed models (GLMMs) as a function of population, gene category, and their interaction at each mutation locus (Supporting Text S1, Eqn 10). To account for unequal gene category sizes, all analyses were repeated using random subsampling across multiple proportions of each gene category. Full methodological details are provided in Supporting Text S1.

### Genome-environment association analyses (GEA) in *M. sylvestris* and overlap with DEGs

To investigate the genomic basis of local climate adaptation in European wild apples, we leveraged its extensive geographic sampling and focused subsequent analyses solely on *M. sylvestris*.

A total of 5.5 million genome-wide SNPs from 64 *M. sylvestris* individuals (Chen et al. 2026)were analysed (Supporting Texts S1 and S2, supplementary table S18). After quality filtering and linkage disequilibrium (LD) pruning using a threshold of 0.2 (PLINK (Purcell et al. 2007)), 26,009 SNPs were retained for population-structure inference with fastStructure (Raj et al. 2014). LD pruning was applied only for population structure analyses; all 5.5 million SNPs were retained for GEA analyses using LFMM (Frichot et al. 2013) and Samβada (Duruz et al. 2019). Six uncorrelated bioclimatic variables (BIO1, BIO2, BIO3, BIO4, BIO8, BIO12) were selected for downstream analyses (Supporting Text S2). Significant climate-associated SNPs were annotated to nearby genes; GEA genes detected by both methods were retained.

Finally, DEGs identified in *M. sylvestris* specific analysis were intersected with (i) genes associated with climate variables identified by GEA (Supporting Text S2) and (ii) outlier genes detected using *D_XY_*. This integrative framework allowed us to identify robust candidate genes underlying local adaptation, supported by concordant signals of regulatory variation, climate-associated genomic variation, and gene divergence.

## Supporting information

Supplementary informations

## Acknowledgements

This research was carried out on the High-Performance Computing resources at New York University Abu Dhabi and on the Core Cluster of the Institut Français de Bioinformatique (IFB) (ANR-11-INBS-0013). We thank Adrien Falce, Olivier Langella, and Benoit Johannet for their help and support at the INRAE Génétique Quantitative et Evolution - Le Moulon lab. We are grateful to Elodie Marchadier for her helpful feedback on the study’s results. The authors would also like to thank the Biological Resource Center “RosePom - Pome Fruits and Roses” and its associated staff, especially the VaDiPom Team from UMR1345 IRHS, INRAE, Agrocampus-Ouest, Université d’Angers, as well as the UE HORTI, INRAE, 2018 Horticulture Experimental Facility (doi:10.15454/1.5573931618268674E12), for their work in maintaining the genetic resources and controlled crosses that provided the *Malus domestica* seeds used in the present study.

## Funding

AMC received funding from the European Agricultural Fund for Rural Development (EAFRD/FEADER) through the LEADER program (Île-de-France region) under grant agreement n° RIDF190219CR0110026; ATIP-CNRS Inserm; IDEEV; Tamkeen under the NYU Abu Dhabi Research Institute grant AD 454; and ANR-21-CE20-0005 (PLEASURE). RD was supported by a PhD fellowship from the French Ministry of Higher Education and Research through the Graduate School ED SEVE 567 (Plant Sciences: From Genes to Ecosystems). AMC and SB were supported by the technical and human resources of the CNRS through the IR ECOTRONS and the CEREEP-Ecotron Île-de-France (CNRS/ENS UMS 3194), including additional funding from the Île-de-France Regional Council under the DIM R2DS program (I-05-098/R, 2011-11017735, 2015-1657) and the French “Investissements d’Avenir” program implemented by the ANR (ANR-10-EQPX-13-01 Planaqua; ANR-11-INBS-0001 AnaEE France). AR and TU received funding through project numbers 88-PHE and BIOCLIMPACT (PN-IV-P8-8.1-PRE-HE-ORG-2024-0223 and PN23020401/7N/03.01.2023) from the MCDI.

## Authors’ contributions

AMC initiated the study; AMC and SB conceived it and secured funding. AMC, AV, CR, ND, SB, SC, AH, AR, TK, TMU, GMV, LF, FL, and GA collected samples and performed the experiments. RD, XC, AMC, JS, ND, MC, and KA analyzed the data. RD, XC, AMC, NC, and SB interpreted the results. AMC, RD, JS, and XC wrote the manuscript with critical input from co-authors. All authors revised and approved the final manuscript.

## Competing interests

The authors declare that they have no competing interests.

## Data Availability

All scripts are available at https://github.com/CornilleEclecticLab/Apple-climate_change_Ecotron. The SSR and SNP VCF data are available on Zenodo: https://doi.org/10.5281/zenodo.18186527. All raw RNA-seq data have been deposited in the NCBI Sequence Read Archive (SRA) under accession PRJNA1399139. All DNA-seq data used in this study have been deposited in the NCBI under BioProject accession number PRJNA1252109.

## Additional files

The following supplementary figures, tables, and methods are provided as Additional file 1 (Supplementary Information), referenced throughout the main text.

**Supplementary fig. S1.** Geographic origin, hybrid ancestry, and genetic differentiation of the reference populations used in this study.

**Supplementary fig. S2.** Genetic relatedness and transcriptomic variation among populations.

**Supplementary fig. S3.** Combined residual trait means from linear mixed models across simulated ecotron climates

**Supplementary fig. S4.** Principal component analysis (PCA) of phenotypic trait variation among apple seedlings.

**Supplementary fig. S5.** Distribution of circadian sampling times relative to sunrise across simulated climate treatments.

**Supplementary fig. S6.** Core climate treatment DEGs overlap with the Armenian climate treatments and without the Armenian climate treatments.

**Supplementary fig. S7.** Gene Ontology enrichment of climate-responsive DEGs identified using a relaxed core-response criterion.

**Supplementary fig. S8.** Distribution and enrichment of core climate-treatment DEGs across WGCNA co-expression modules.

**Supplementary fig. S9.** Gene Ontology (GO) term enrichment analysis across WGCNA modules enriched with core climate-treatment differentially expressed genes (DEGs).

**Supplementary fig. S10**. Correlation between the expression levels of genes within each WGCNA co-expression module and early phenotypic traits in apple seedlings.

**Supplementary fig. S11.** Pairwise Pearson’s correlation coefficients among 55 environmental variables from WorldClim2.

**Supplementary fig. S12.** Manhattan plot of genotype × environment associations (GEAs) in *Malus sylvestris* across six bioclimatic variables based on statistics from LFMM2.

**Supplementary fig. S13.** Manhattan plot of genotype × environment associations (GEAs) in *Malus sylvestris* based on G-score statistics from Samβada.

**Supplementary fig. S14.** Bioinformatic pipeline for SNP detection and estimation of nucleotide diversity (π), absolute genetic divergence (D_XY_), and genetic differentiation (F_ST_) from RNA-seq data.

**Supplementary table S1.** Seedling provenance and experimental use across climate simulations.

**Supplementary table S2.** Seedling survival across ecotron climate simulations and statistical associated tests.

**Supplementary table S3.** Number of seedlings measured for each trait across genetic groups/populations, climate simulations, and experimental years.

**Supplementary table S4.** Climate-induced plasticity across traits and genetic groups/populations from random-slope regression mixed models (RRMMs).

**Supplementary table S5.** Gene expression profiles of genes expressed in this study.

**Supplementary table S6.** Core climate-treatment genes with known roles in flowering regulation.

**Supplementary table S7.** Results of Fisher’s Exact Tests for *D_XY_* outlier enrichment and depletion across DEG categories.

**Supplementary table S8.** Linear mixed model results for nucleotide diversity statistics obtained from SNP calling of RNA-seq data.

**Supplementary table S9.** Estimated marginal means and pairwise contrasts of log-transformed π*N*/π*S* ratios across gene categories.

**Supplementary table S10.** Estimated marginal means and pairwise contrasts of log-transformed diversity ratios between apple genetic groups/populations across two gene categories, Background genes and dom-wild DEGs.

**Supplementary table S11.** Estimated marginal means and pairwise contrasts of log-transformed diversity ratios between domesticated and wild apple genetic groups/populations.

**Supplementary table S12.** Binomial generalized linear mixed model results for normalized deleterious mutation burden by genotype state at mutation loci measured in the dataset from Chen et al. 2026.

**Supplementary table S13.** Estimated marginal means and pairwise contrasts of normalized deleterious mutation burden by population, gene category, and genotype state.

**Supplementary table S14.** Genes presenting SNPs with a significant gene × environment association.

**Supplementary table S15.** Genes presenting climate-aligned expression in *Malus sylvestris* only expression test.

**Supplementary table S16.** Year effect on phenotypic traits for apple seedlings in the ecotron experiment.

**Supplementary table S17.** Best linear unbiased predictors (BLUPs) of year effects on phenotypic traits in the ecotron experiment.

**Supplementary table S18.** Sample information and geographic origin of the *Malus sylvestris* used in the GEA analysis.

**Supplementary table S19.** Nucleotide diversity across populations and functional gene categories.

## Notes

https://doi.org/10.5281/zenodo.18186528

### Competing Interest Statement

The authors have declared no competing interest.

### Summary of Updates

- Format was changed to Introduction, Results, Discussion, Methods - Introduction was modified. - Additional precision was given to the methods section and supporting material. - Added supplementary figures on Hybrid index. - Added supplementary tables on Pi statistics. - Reorganisation of supplementary tables and figures to follow the new format. - Modifications to the deleterious mutations analysis were performed to follow the framework from Chen et al. 2026.

