## Supplementary informations for "Conserved transcriptional plasticity, not local adaptation, dominates early climate responses across wild and cultivated apples"

The following Supporting Information is available for this article:

**Supplementary fig. S1.** Geographic origin, hybrid ancestry, and genetic differentiation of the reference populations used in this study.

**Supplementary fig. S2.** Genetic relatedness and transcriptomic variation among populations.

**Supplementary table S19.** Nucleotide diversity across populations and functional gene categories.

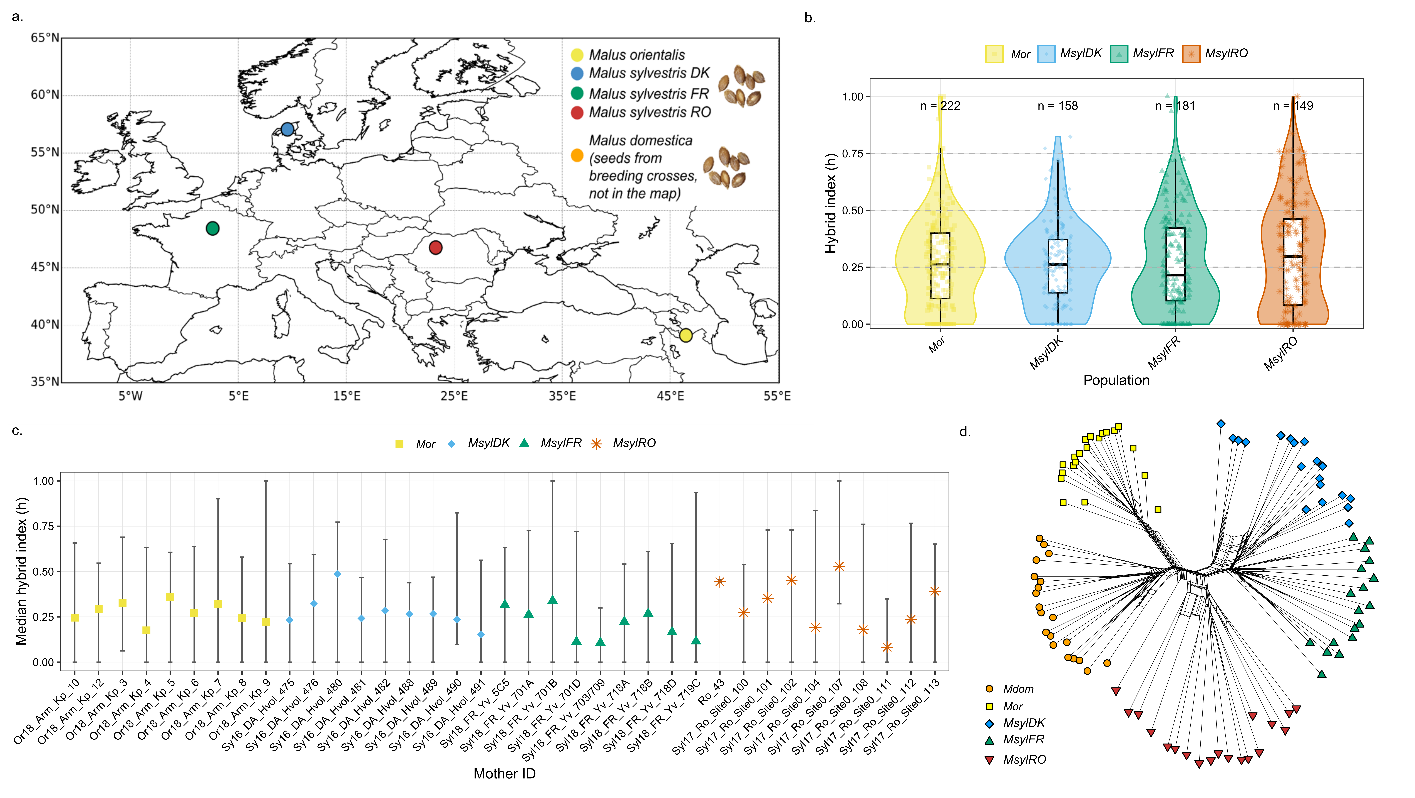
 **Supplementary fig. S1. Geographic origin, hybrid ancestry, and genetic differentiation of the reference populations used in this study.** (a) Geographic origin of the wild reference populations used to estimate the hybrid index (h): *Malus orientalis* from Armenia and *M. sylvestris* from Denmark, France, and Romania. Reference *M. domestica* genotypes originated from breeding accessions described in Cornille et al. (2015) and are therefore not represented on the map. (b) Distribution of the hybrid index (*Pdom*) for seedlings from each wild population. Violin plots show the density of individual values, boxplots indicate the median and interquartile range, and points represent individual seedlings. Sample sizes are indicated above each violin. A value of *Pdom* = 1 corresponds to pure *M. domestica* ancestry, whereas *Pdom* = 0 corresponds to pure wild ancestry (*M. sylvestris* or *M. orientalis*). (c) Median hybrid index and observed range for each maternal family, illustrating the variation in domesticated ancestry among offspring from different mother trees. For seedlings that could not be genotyped, *Pdom* values were imputed as the mean hybrid index of successfully genotyped siblings from the same maternal family. (d) SplitsTree network inferred from genome-wide SNPs called from RNA sequencing reads showing the genetic relationships among the reference populations. The network reveals clear genetic differentiation among the four wild populations, supporting their classification as distinct genetic groups despite the limited resolution of the microsatellite dataset.

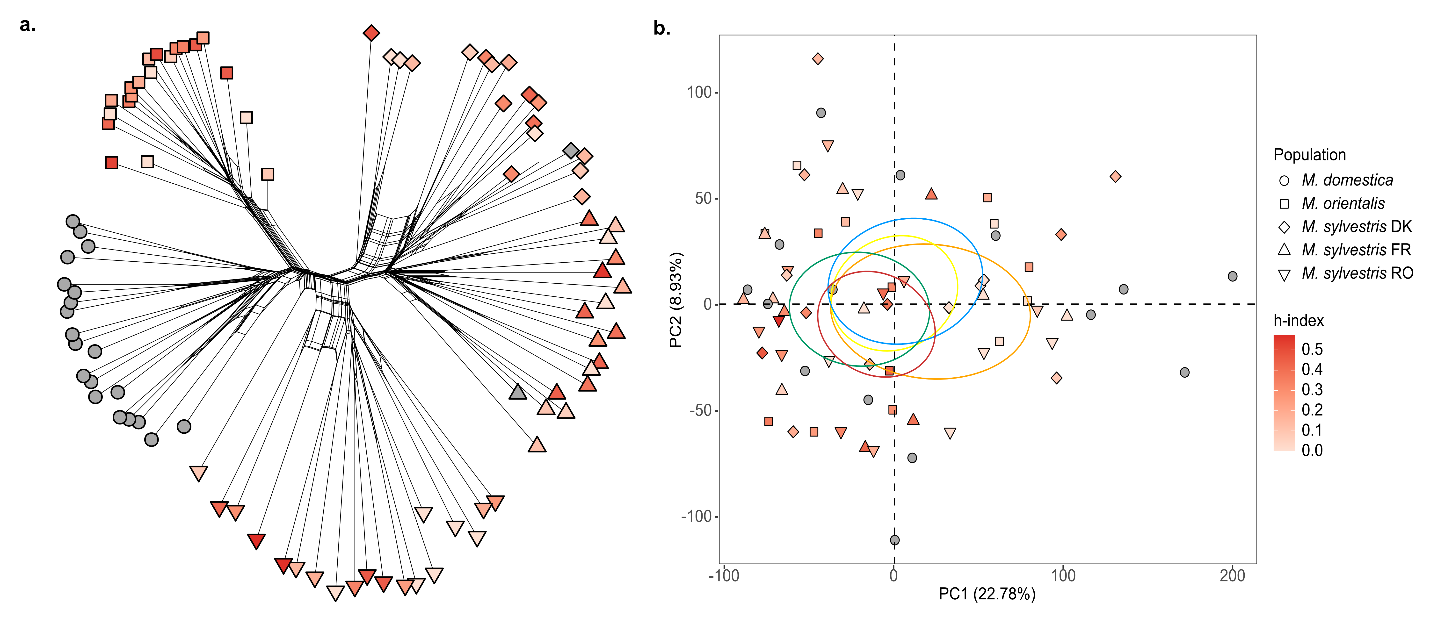

**Supplementary fig. S2. Genetic relatedness and transcriptomic variation among populations.** (a) A Neighbor-Net split network illustrating the genetic relationships among individual samples based on bitwise distances computed from genomic variant data. Tip shapes correspond to the distinct apple populations. The node fill color represents the h-index, mapped on a continuous scale from light pink/white to dark red, with dark grey nodes indicating missing h-index values (NA). (b) Principal Component Analysis (PCA) of normalized RNA-seq gene expression counts. The scatter plot displays the projection of individual samples onto the first two principal components (PC1 and PC2). Sample shapes denote the populations (*M. domestica*, *M. orientalis*, and *M. sylvestris)* from Denmark [DK], France [FR], and Romania [RO]. Similar to panel a, the fill color of the symbols indicates the individual's h-index. The colored ellipses represent the barycentric confidence ellipses computed for each respective population

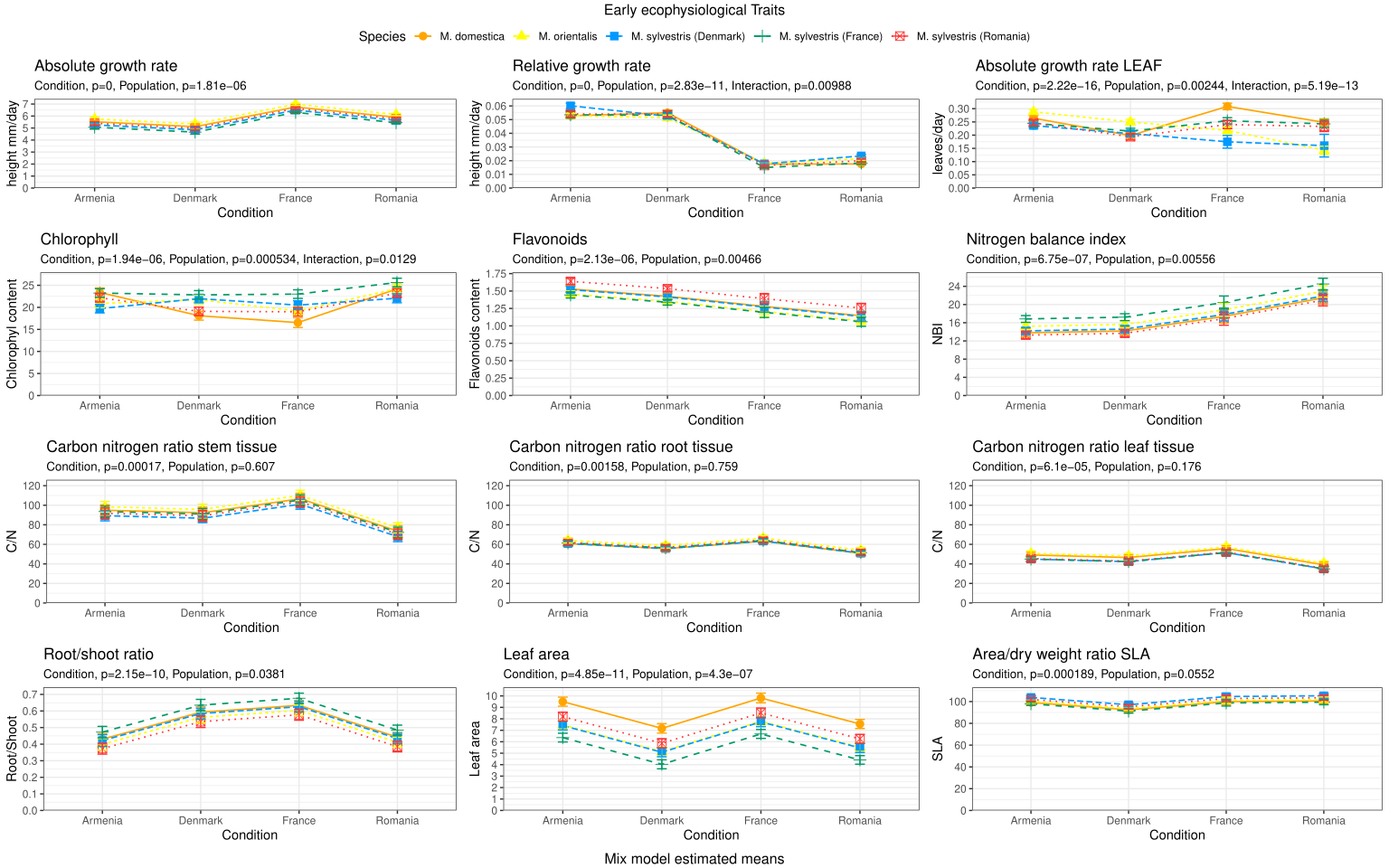

### **Supplementary fig. S3. Combined residual trait means from linear mixed models across simulated ecotron climates.** Plots show residual means (best linear unbiased predictions [BLUPs]) for physiological traits measured under four climate treatments: Armenia and Denmark (in 2019) and France and Romania (in 2021). Residuals were extracted from linear mixed models correcting for population, climate, and year effects. Colors indicate genetic groups/populations: *Malus domestica* (yellow), *M. orientalis* (orange), and *M. sylvestris* originating from Denmark (blue), France (green), or Romania (red), respectively.

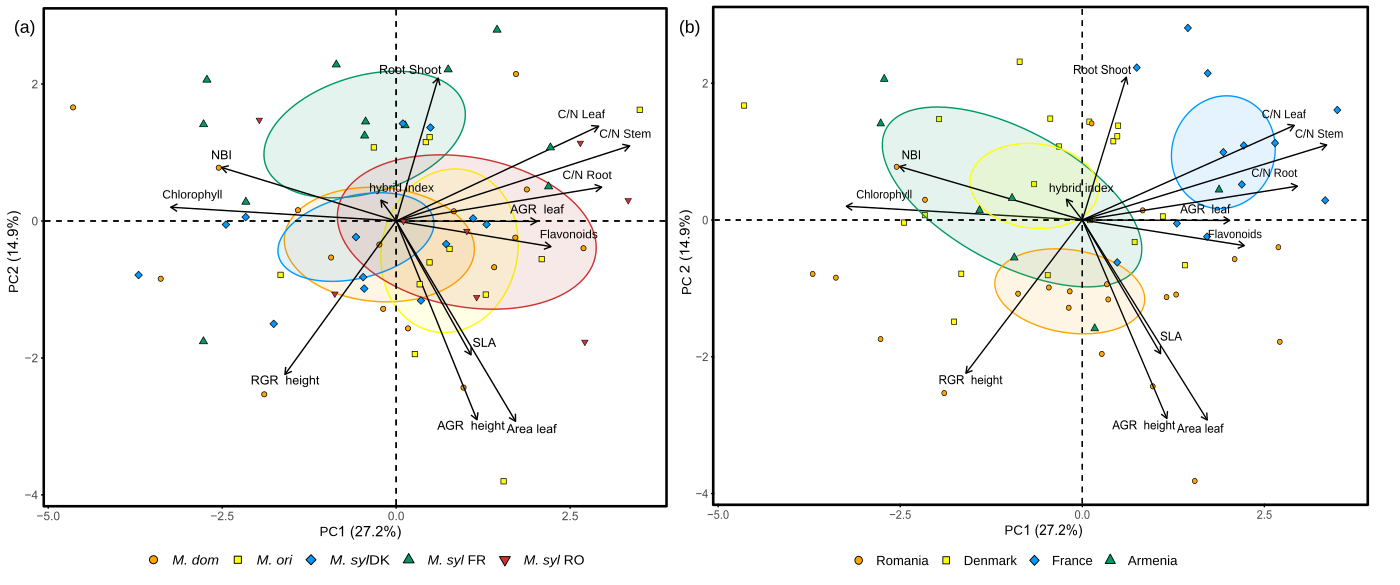

### **Supplementary fig. S4. Principal component analysis (PCA) of phenotypic trait variation among apple seedlings.** (a) PCA plot illustrating phenotypic differentiation among seedlings from five genetic groups/populations: *Malus domestica*, *M. orientalis*, and *M. sylvestris* (from Denmark, France, or Romania) **(b)** PCA plot showing phenotypic variation among seedlings across four simulated ecotron climates: Armenia and Denmark (2019) and France and Romania (2021). Percentages given along each axis indicate the proportion of total phenotypic variance explained by each principal component. Clustering patterns reflect the combined effects of population origin and climate treatment on early seedling trait expression.

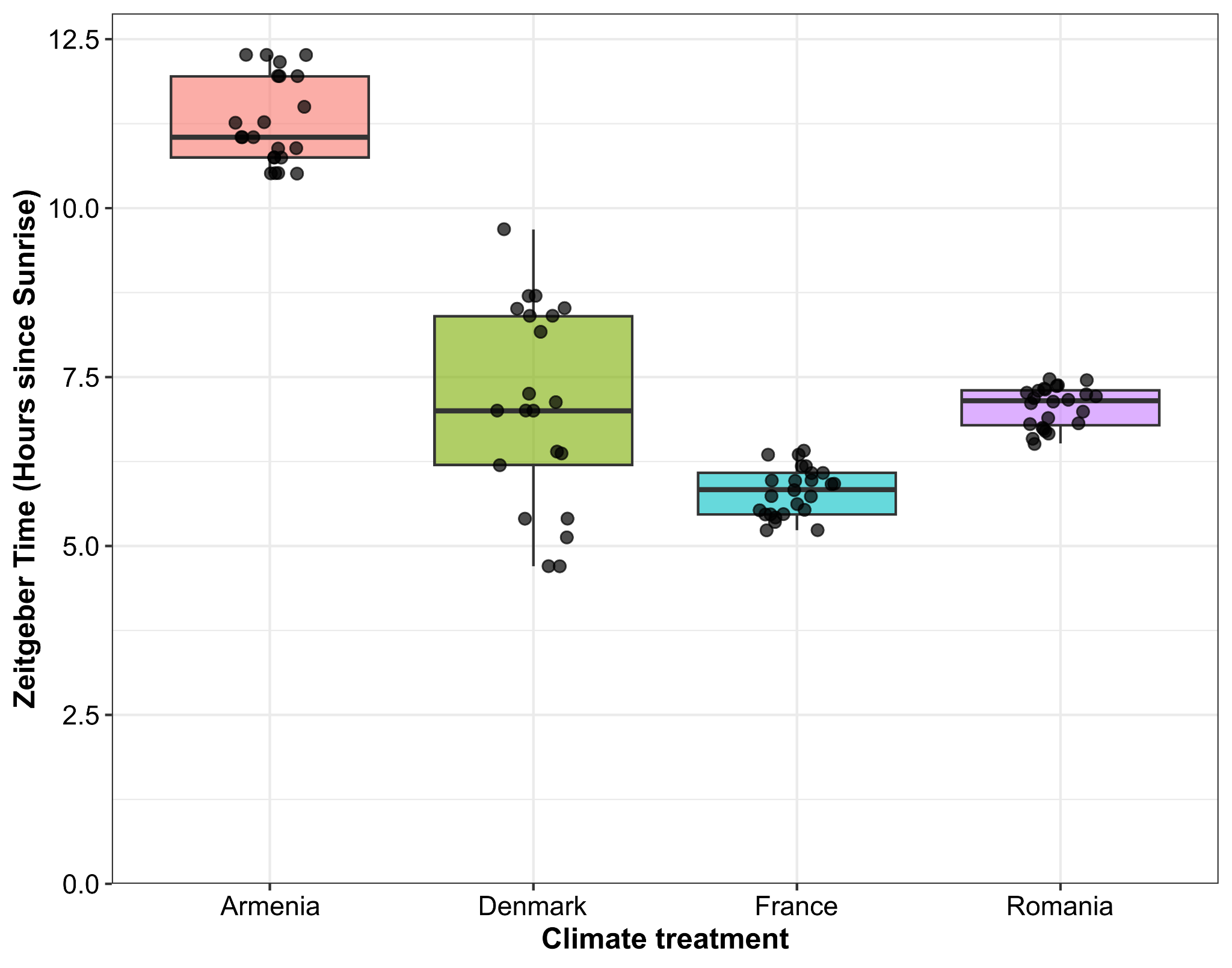

### **Supplementary fig. S5. Distribution of circadian sampling times relative to sunrise across simulated climate treatments.** Boxplot showing the variation in Zeitgeber Time (ZT) for samples collected across four simulated ecotron climates: Armenia and Denmark (2019) and France and Romania (2021). The y-axis displays the sampling time in hours since sunrise, with ZT 0 marking sunrise for each condition. Individual sampling times are represented as overlaid points to illustrate the raw data distribution. Variations in ZT distributions reflect the differences in localized photoperiods and experimental sampling schedules across the simulated ecotron environments.

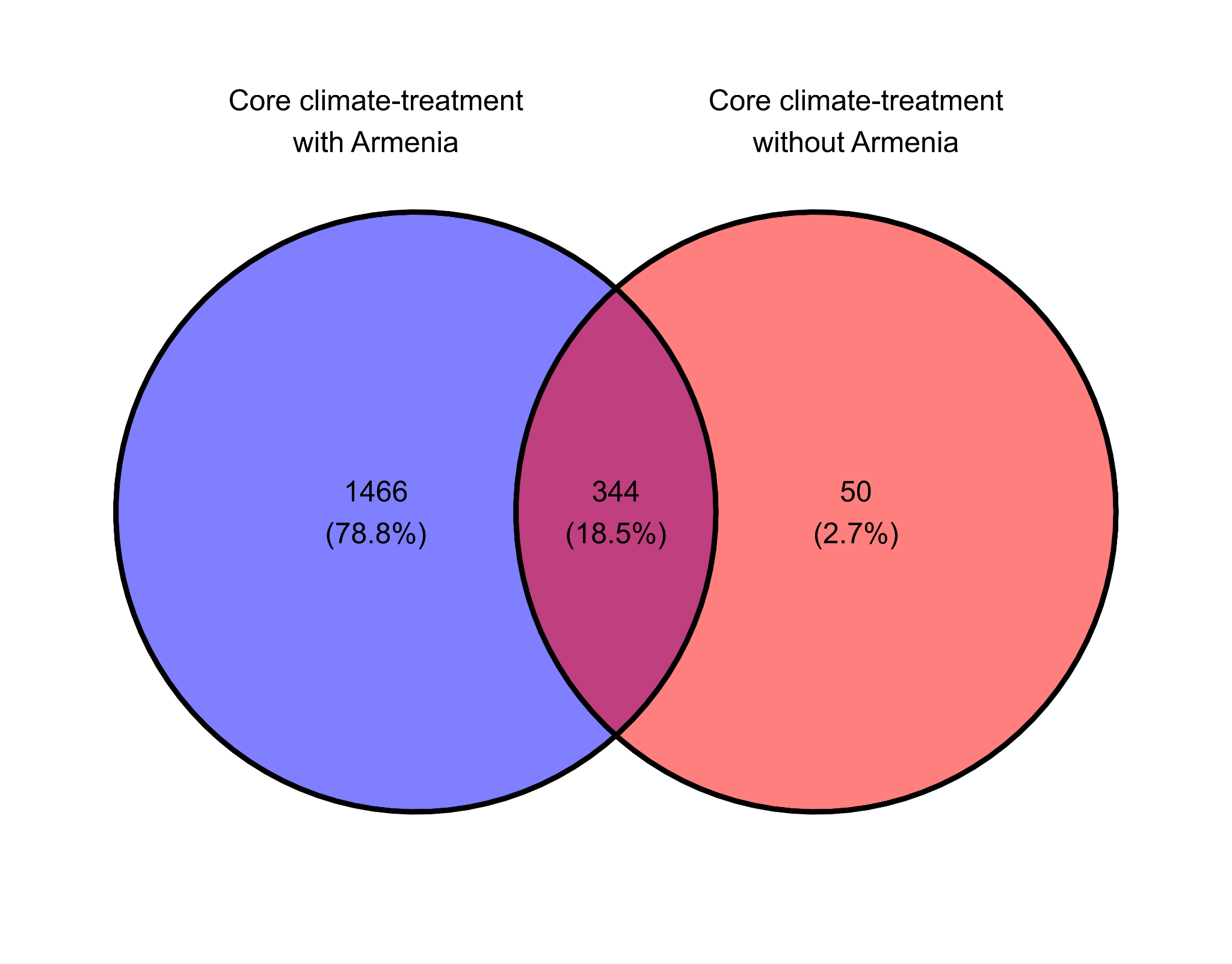

**Fig S6. Overlap between core climate-treatment DEGs identified with and without the Armenian treatment.** Venn diagram showing the overlap between core climate-treatment differentially expressed genes (DEGs) identified using all four climate treatments (including Armenia) and those identified after excluding the Armenian treatment (France, Romania, and Denmark only). Core climate-treatment DEGs are genes that are consistently responsive to climate treatments across all sampled lineages. Numbers indicate the number of genes in each category, with percentages calculated relative to the total number of DEGs identified across both analyses.

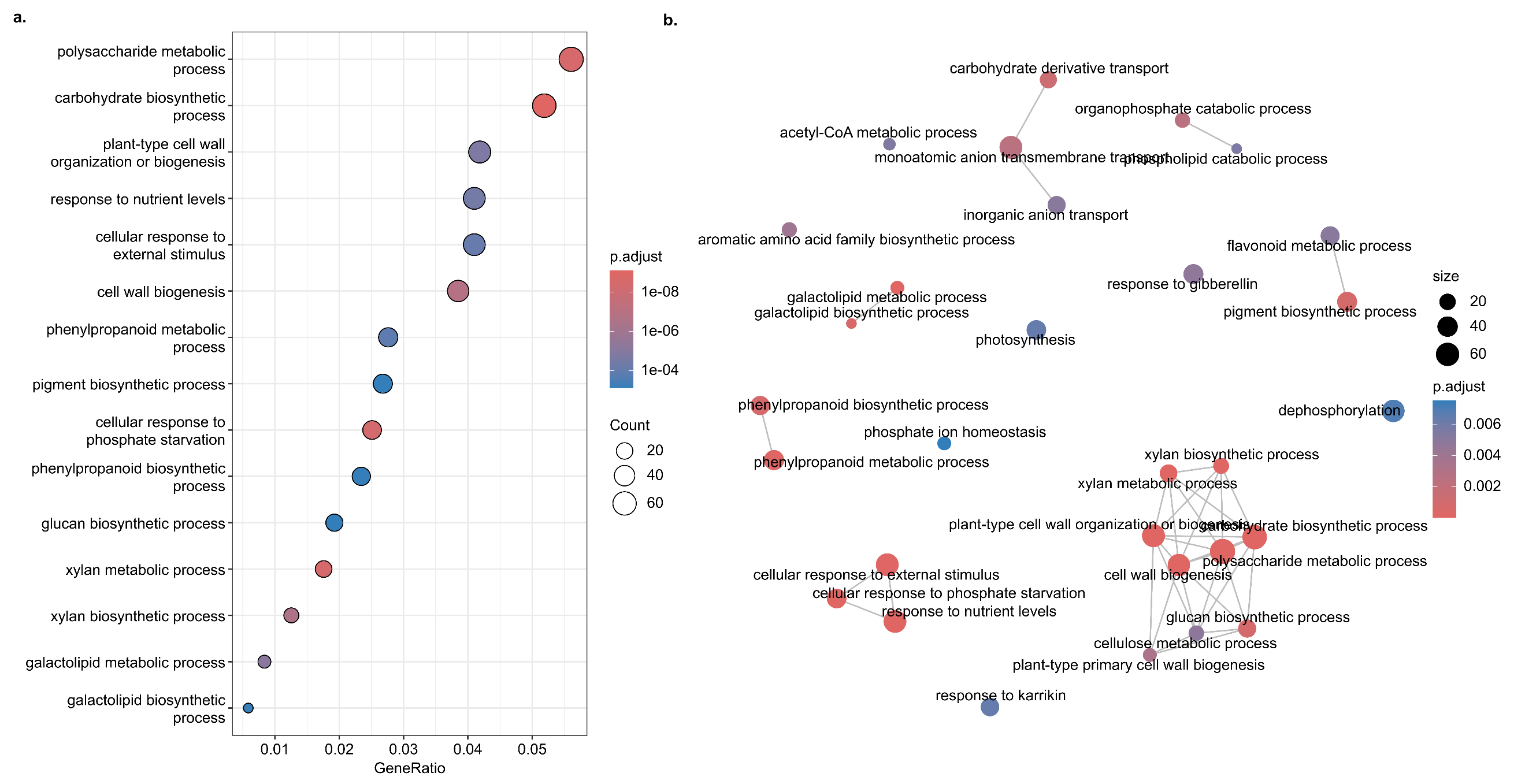

**Supplementary fig. S7. Gene Ontology enrichment of climate-responsive DEGs identified using a relaxed core-response criterion.** GO enrichment analysis of the 2,372 genes showing a significant climate response in at least two of the five populations, based on the three-treatment dataset excluding the Armenian treatment. **(a)** Dot plot of significantly enriched GO terms. Dot size represents the number of genes associated with each GO term, the x-axis represents the GeneRatio, and color indicates the Benjamini–Hochberg-adjusted *P*-value. **(b)** Network representation of enriched GO terms based on semantic similarity. Nodes represent enriched GO terms, with node size proportional to the number of associated genes and color indicating the adjusted *P*-value; edges connect semantically related GO terms.

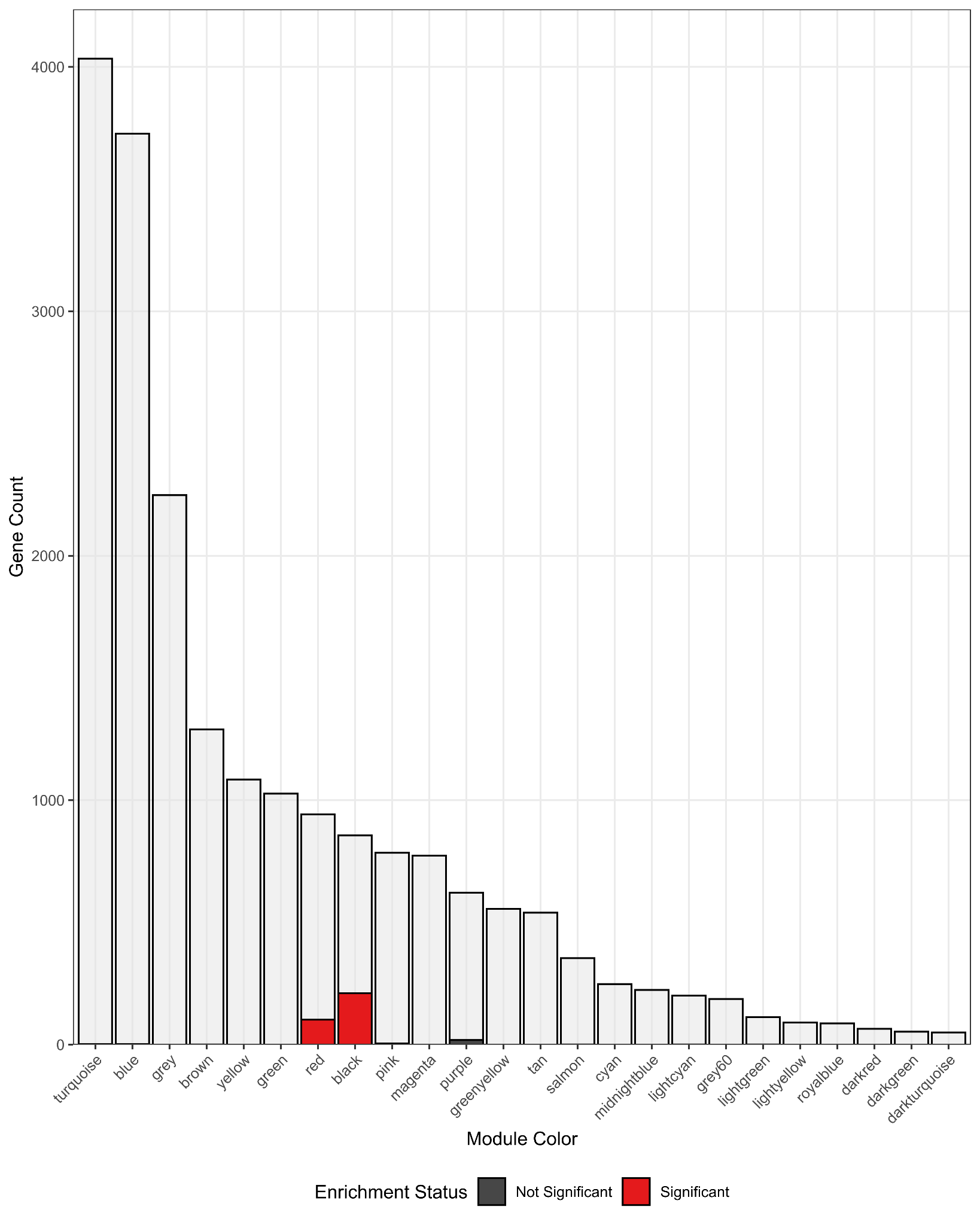

**Supplementary fig. S8**. **Distribution and enrichment of core-climate treatment DEGs across WGCNA co-expression modules.** The bar plot illustrates the overlap between co-expression modules identified via Weighted Gene Co-expression Network Analysis (WGCNA) and core climate-treatment DEGs (n=394). For each module (x-axis), the light grey background bars represent the total number of genes assigned to that cluster, while the foreground bars indicate the number of core climate genes contained within. Statistical significance was assessed using a one-sided Fisher’s Exact Test to identify modules with a higher-than-expected proportion of core climate treatment DEGs relative to the genomic background (N=20,157). p-values were adjusted for multiple testing using the Benjamini-Hochberg (FDR) method. Modules colored in red represent significant enrichment (FDR < 0.05), while modules in dark grey show no significant over-representation. The "grey" module denotes genes not assigned to any specific co-expression cluster.

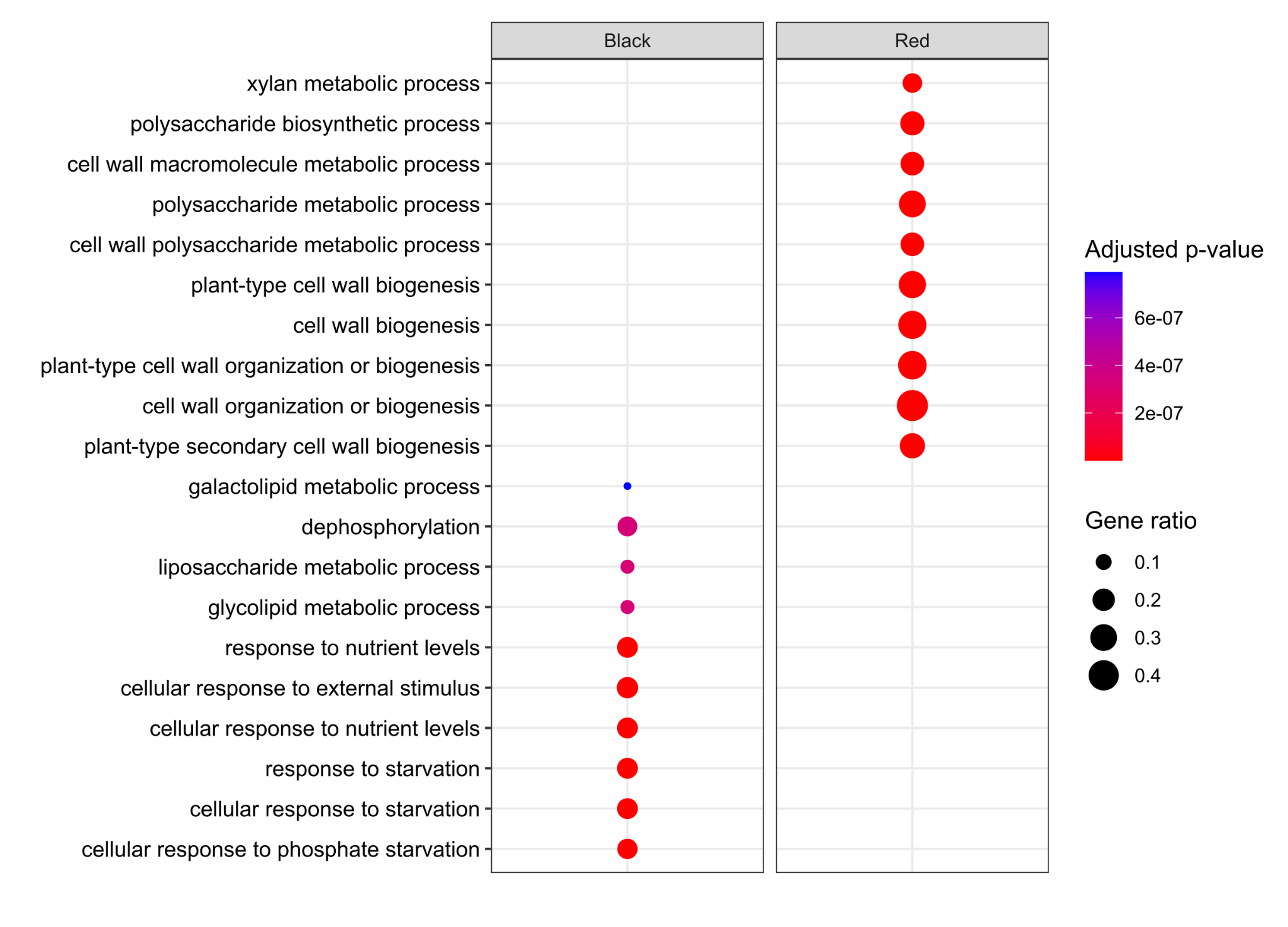

### **Supplementary fig. S9. Gene ontology (GO) term enrichment analysis across WGCNA modules enriched with core-climate treatment differentially expressed genes (DEGs).** GO enrichment was assessed within each module, and only modules significantly enriched with climate-responsive expression patterns were included. The colors at the bottom correspond to the WGCNA assignment.

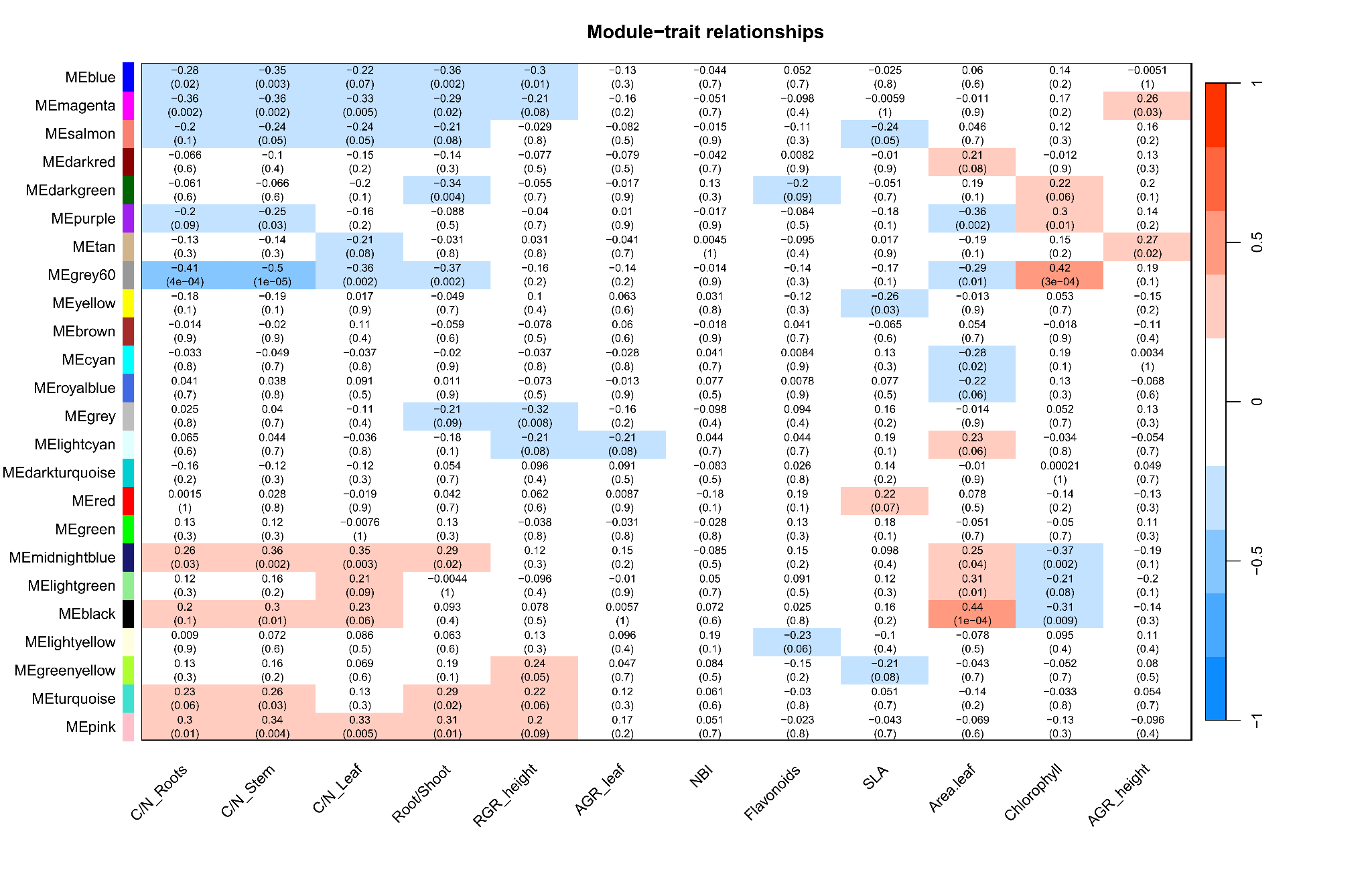

### **Supplementary fig. S10**. **Correlation between the expression levels of genes within each WGCNA co-expression module and early phenotypic traits in apple seedlings.** The correlation matrix shows the relationship between the expression levels of co-expressed modules (color-coded by WGCNA) and early-stage phenotypic traits measured in the ecotron experiment. Analyses were conducted on seedlings from *M. domestica* (cultivated apple), *M. orientalis* (Caucasian wild apple), and *M. sylvestris* (European crabapple) genetic groups/populations from Denmark, France, and Romania. Each cell represents the correlation coefficient between a co-expression module and a phenotypic trait, with associated significance levels given in parentheses.

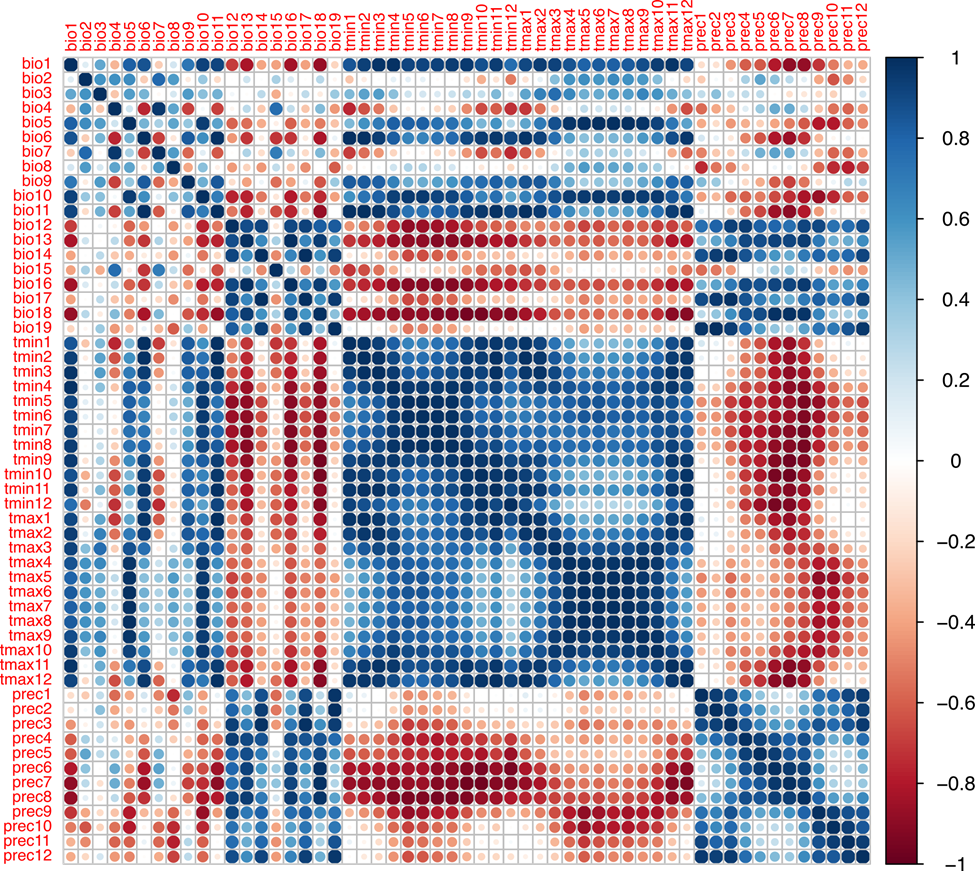

### **Supplementary fig. S11**. **Pairwise Pearson’s correlation coefficients among 55 environmental variables from WorldClim2.** Matrix of pairwise Pearson’s correlation coefficients computed between 55 bioclimatic and environmental variables from the WorldClim2 database. Correlations were calculated using the prepareEnv function from the Rsambada R package. Strong correlations (positive or negative) indicate potential multicollinearity and were used to inform variable selection for downstream genotype×environment association (GEA) analysis.

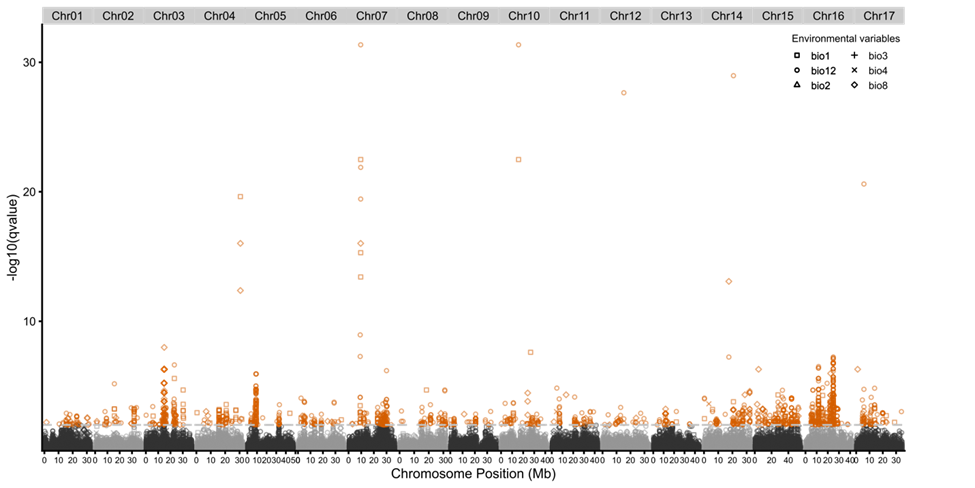

### **Supplementary fig. S12.** **Manhattan plot of genotype×environment associations (GEAs) in *Malus sylvestris* across six bioclimatic variables based on statistics from LFMM2.** The Manhattan plot shows −log₁₀-transformed *q*-values for single-nucleotide polymorphism (SNP) associations across the *M. sylvestris* genome with six selected environmental variables, as identified by LFMM2. Each point represents a SNP tested for association; red circles indicate significant associations with *q*-value < 0.01. Variables analyzed were BIO1 (annual mean temperature), BIO2 (mean diurnal range: mean of monthly [max temp − min temp]), BIO3 (isothermality: BIO2 / BIO7 × 100), BIO4 (temperature seasonality: standard deviation × 100), BIO8 (mean temperature of wettest quarter), and BIO12 (annual precipitation). SNP positions are plotted by genomic location, highlighting loci potentially involved in local adaptation.

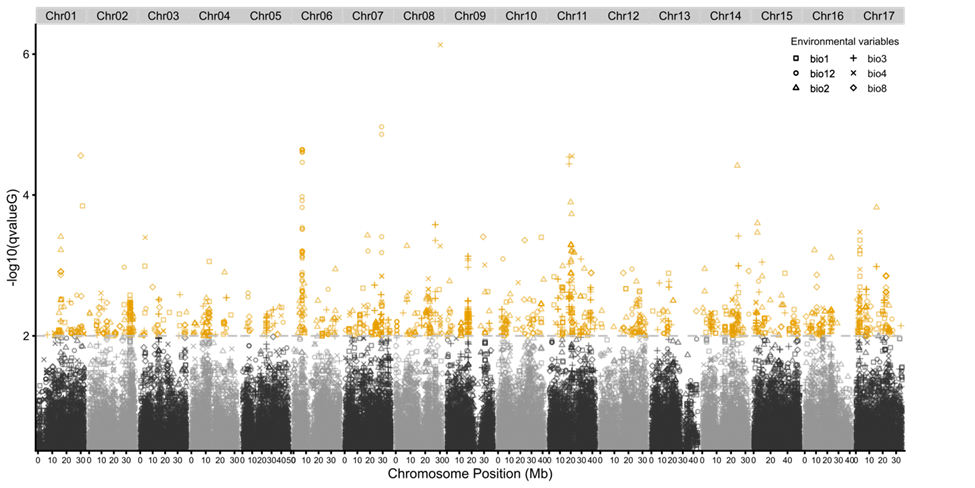

### **Supplementary fig. S13. Manhattan plot of genotype×environment associations (GEAs) in *Malus sylvestris* based on G-score statistics from Samβada.** The Manhattan plot shows −log₁₀-transformed *q*-values for single-nucleotide polymorphism (SNP) associations across the *M. sylvestris* genome with six environmental variables, as calculated using the G-score test in the Samβada framework. Each point represents a SNP tested for association; orange circles indicate statistically significant associations with q-value < 0.01. Variables were BIO1 (annual mean temperature), BIO2 (mean diurnal range: mean of monthly [max temp − min temp]), BIO3 (isothermality: BIO2 / BIO7 × 100), BIO4 (temperature seasonality: standard deviation × 100), BIO8 (mean temperature of wettest quarter), and BIO12 (annual precipitation). SNPs are plotted by chromosomal position to identify genomic regions that may be under local environmental selection.

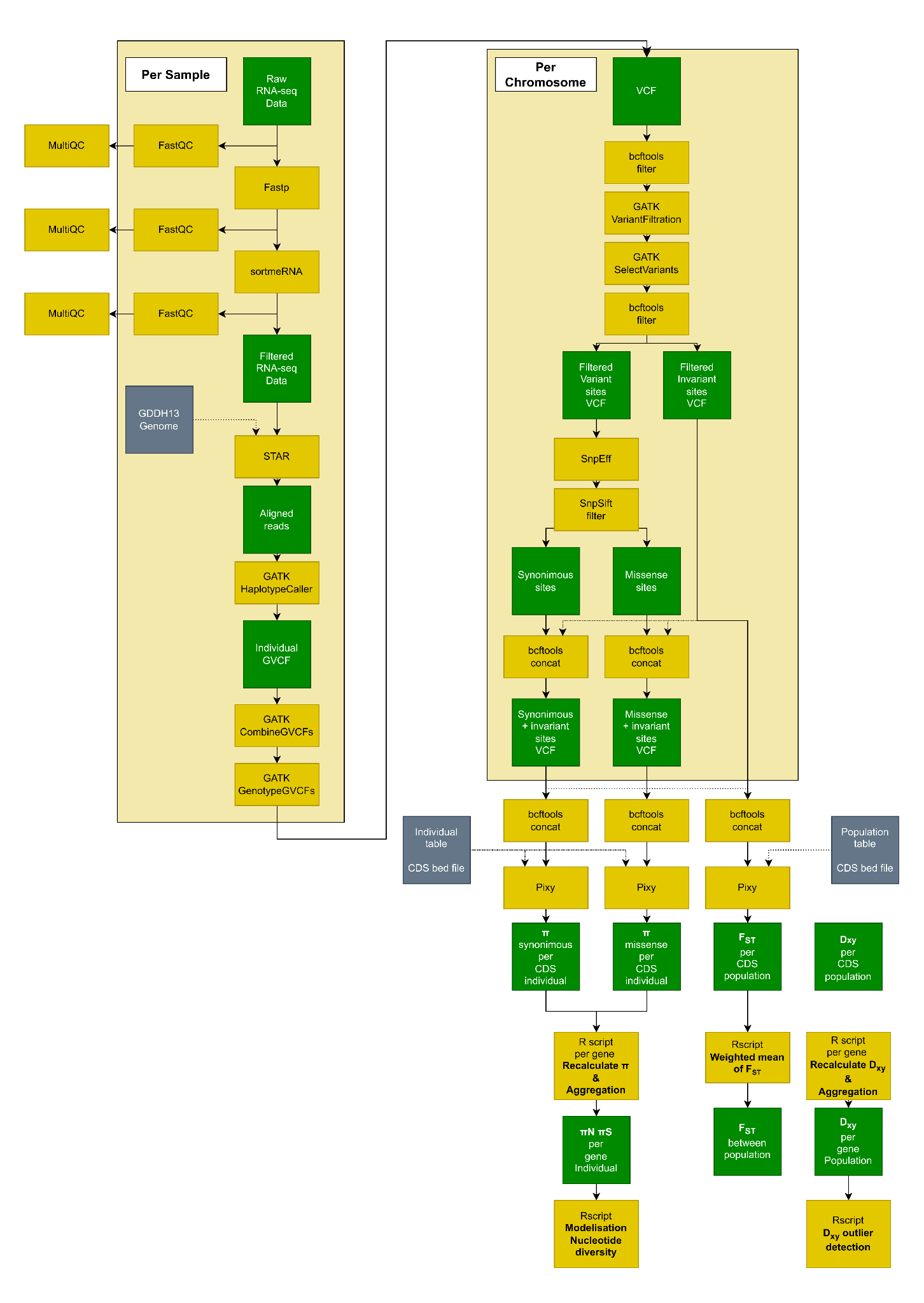

**Supplementary fig. S14.** **Bioinformatic pipeline for SNP detection and estimation of nucleotide diversity (π), absolute genetic divergence (*D_XY_*), and genetic differentiation (*F_ST_*) from RNA-seq data.** Raw RNA-seq reads underwent quality control and filtering, including adapter trimming and removal of low-quality reads, and were then subjected to ribosomal RNA depletion. Filtered reads were aligned to the reference genome, and variants were called using the GATK pipeline. SNPs were subsequently filtered to retain high-confidence variants and annotated prior to downstream analyses. Population genomic statistics, including nucleotide diversity (π) and pairwise *F_ST_*, were then calculated using filtered SNP datasets. This workflow follows standard best practices for RNA-seq-based SNP calling and allows estimation of genetic diversity and differentiation from transcriptome-derived polymorphisms.
