## Supplementary text 1 for "Conserved transcriptional plasticity, not local adaptation, dominates early climate responses across wild and cultivated apples"

### Text S1. Detailed Materials and Methods

Seeds from five apple populations were used: *Malus sylvestris* from Romania, France, and Denmark (Cornille et al. 2015); *M. orientalis* from Armenia, known for its stress tolerance (Volk et al. 2015); and cultivated apple (*M. domestica*) from controlled crosses between cultivars representing a large diversity from traditional or recent compartments maintained by the Biological Resource Center “RosePom - Pome Fruits and Roses” (supplementary table S1).

The experimental material comprised seeds from 15 mother trees for *M. domestica*, 9 for *M. orientalis*, and 9, 9, and 10 mother trees for the Danish, French, and Romanian M. sylvestris populations, respectively (supplementary table S1). Offspring from each mother tree were distributed across the four climate treatments whenever sufficient seeds were available, with the number of seedlings per maternal family and treatment varying according to seed availability (supplementary table S1). In total, the experiment included 258 *M. domestica*, 251 *M. orientalis*, 241 Danish *M. sylvestris*, 252 French *M. sylvestris*, and 247 Romanian *M. sylvestris* seedlings (N = 1,249). Seeds per mother tree were surface-sterilized (0.5% [v/v] bleach for 20 min) and vernalized in 180-ml tubes containing a damp mixture of sand and vermiculite in equal volumes for 3 months at 4°C in darkness. Seeds from the same mother trees were used across all climate treatments, with some exceptions owing to limited seed availability (supplementary table S1).

Four climate treatments were simulated in the ecotron growth chamber facilities at CNRS (Foljuif, France) (Roy et al. 2021). These climate chambers allow precise control over environmental variables, closely mimicking natural field conditions. Owing to spatial and temporal constraints, the experiment was divided over two years: two climate treatments (Denmark and Armenia) were simulated in 2019, and two others (Romania and France) in 2021 (Fig. 1). Year effects were accounted for in all subsequent statistical analyses.

Each climate treatment corresponded to the local temperature and humidity profiles of meteorological stations located near the seed collection sites: France (longitude: 2.63, latitude: 48.44), Denmark (longitude: 9.55, latitude: 57.06), Romania (longitude: 23.3, latitude: 46.78), and Armenia (longitude: 46.47, latitude: 39.13), as well as the day length.

To expose the seeds to the historical climate conditions of the four native sites, they were placed in ECOTRON climate chambers programmed to replicate site-specific temperature and humidity profiles from May 15th to July 15th. The chambers were programmed to adjust to specific temperature and humidity setpoints (targets) every three hours. These precise setpoints were calculated by averaging 30 years of historical data (1985-2014) over 3-hour intervals for the same seasonal window (May 15-July 15). A common calendar window was deliberately used for all four climate simulations to provide the same temporal reference across treatments and populations. The objective was to compare responses to the climatic conditions experienced at each site during this fixed early-growing-season period, rather than to reproduce site-specific phenological onset. This process yielded a total of 488 distinct climate targets per native site, enabling the chambers to accurately simulate natural diurnal and seasonal fluctuations. The historical temperature and humidity datasets used to generate these targets were retrieved from the Climate Data Interface (CDI). Raw data were extracted in NetCDF format and subsequently converted to CSV format for downstream processing using the R package ncdf4 (Pierce 2025).

In addition to temperature and humidity controls, daylight periods were adjusted daily for each chamber to replicate the natural photoperiod variation observed between May 15th and July 15th at each native site. To achieve this, precise daily daylength data for each geographic location were extracted from the ptaff database and incorporated into the chamber programming accordingly (Fig. 1).

Following stratification, seeds were sown into Jiffy pellets arranged in 20-hole trays and placed in the ecotron chambers. Sowing dates were July 17, 2019, for the Armenian and Danish conditions (309 and 306 seeds, respectively; supplementary table S2), and February 17, 2021, for the Romanian and French conditions (317 and 316 seeds, respectively; supplementary table S2). Each 20-hole tray was rotated daily within the chamber to minimize micro-environmental variation, and seedlings were watered weekly (1 L per 20-hole tray, evenly distributed). Seedlings were grown for 2 months: from July 17 to September 18, 2019, and from February 17 to April 21, 2021.

Wild seedlings were genotyped using 12 microsatellite (SSR) markers to quantify the extent of introgression from *M. domestica*. These genotypes were subsequently used to estimate a hybrid index (*Pdom*) for each seedling, as detailed below. Genomic DNA was extracted using a NucleoSpin Plant DNA Extraction Kit II (Macherey-Nagel, Düren, Germany) according to the manufacturer’s instructions. Microsatellites were amplified using a Multiplex PCR Kit (QIAGEN, Inc.) and a multiplex PCR protocol. Twelve microsatellite markers were used, namely Ch01f02, Ch01f03, Ch01h01, Ch01h10, Ch02c06, Ch02c09, Ch02c11, Ch02d08, Ch03d07, Ch04c07, Ch05f06, GD12, and Hi02c07 in four multiplex reactions (MP01-MP04; (Cornille et al. 2012)). PCR was performed in a final reaction volume of 15 μL (7.5 μL of QIAGEN Multiplex Master Mix, 10-20 μM of each primer, with the forward primer labeled with a fluorescent dye, and 10 ng of template DNA). A touchdown PCR program was used (initial annealing temperature of 60°C, decreasing by 1°C per cycle to 55°C). Genotyping was performed at the GENTYANE platform (INRAE Clermont-Ferrand) on an ABI PRISM X3730XL, with 2 µl of GS500LIZ size standard (Applied Biosystems). Alleles were scored with GENEMAPPER 4.0 software (Applied Biosystems). Only multilocus genotypes with less than 10% missing data were retained for analysis.

For each individual, a hybrid index (*Pdom*) was computed to represent the proportion of alleles assigned to the *M. domestica* gene pool using the introgress package in R (Gompert and Buerkle 2010). Reference genotypes for *M. domestica* and *M. sylvestris* (France, Romania, Denmark) were taken from (Cornille et al. 2015), while reference genotypes for *M. orientalis* were obtained from (Bina et al. 2022). A value of *Pdom* = 1 corresponds to pure *M. domestica* genotypes, whereas *Pdom* = 0 corresponds to pure wild genotypes (*M. sylvestris* or *M. orientalis*). All *M. domestica* seedlings were assumed to have *Pdom* = 1. For seedlings for which DNA extraction failed, *Pdom* values were imputed as the average hybrid index of successfully genotyped siblings from the same mother tree.

##### Growth and physiological traits

Seedling survival, growth, photosynthetic activity, and carbon-related traits were measured throughout the experiment. Sample sizes varied owing to time constraints and destructive sampling at the end of the experimental period.

All seedlings were monitored every 2-3 days to record germination time, seedling height, and number of leaves. These data were used to calculate growth dynamics: absolute growth rate (AGR), following (Radford 1967), as AGR = Δtrait / Δdays, and relative growth rate (RGR), following (Briggs et al. 1920), as RGR = [ln(final trait value) − ln(initial trait value)] / Δtime. AGR and RGR were calculated for height and leaf number. For each seedling, the final AGR was defined as the mean AGR across the growth period, excluding values before germination or after death.

Photosynthetic traits were measured on day 61 in a consistent 2-h window to avoid diurnal variation, using a subsample of 15 seedlings per population and treatment (*N* = 260; supplementary tables S1, S3). The subsample included at least five mother trees per population (Table S1), with three seedlings per mother tree whenever possible (supplementary table S1). Chlorophyll content (Chl), flavonoid content (Flav), and nitrogen balance index (NBI) were measured from three leaves per seedling using a portable Dualex device (Force-A, Orsay, France). Chl reflects the concentration of photosynthetic pigments (µg/cm²), Flav indicates the accumulation of phenolic compounds associated with stress responses and ultraviolet (UV) light protection, and NBI (Chl/Flav) serves as a proxy for nitrogen-to-carbon allocation in leaves.

At the end of the experiment, destructive sampling of 5-19 seedlings per group allowed measurements of Carbon and nitrogen content determined by dry combustion using a CN auto-analyzer (Flash EA 1112 series, Thermo Finnigan). The same individuals were used to assess morphological traits: specific leaf area (SLA; leaf area/dry mass) and root-to-shoot ratio (root dry mass/aboveground dry mass).

In total, 12 traits grouped into four categories were analyzed, with each trait reflecting a different sampling scale: growth traits (AGR height, AGR leaf, RGR height), measured on all seedlings; photosynthetic traits (Chl, Flav, NBI), measured on 15 seedlings per population and treatment; tissue stoichiometry traits (C/N leaf, C/N stem, C/N root), which required destructive sampling; and morphological traits (SLA, root/shoot, leaf area), measured on the same destructively sampled individuals. Whenever possible, the same seedlings were used across trait categories to ensure comparability (supplementary tables S1, S3).

#### Survival analyses

#### Survival was scored as a binary trait (alive = 1, dead = 0) on day 61 after transplantation and analyzed separately from the twelve phenotypic traits (supplementary table S2a), which were measured only on surviving seedlings. In contrast, survival analyses included both surviving and non-surviving individuals. Analyses focused on five populations: *M. domestica* (MD), *M. orientalis* (MOr), and three *M. sylvestris* populations originating from France (MSylFR), Denmark (MSylDK), and Romania (MSylRO) (supplementary table S2).

#### To test for population, climate treatment, and population × climate treatment effects on survival, we fitted binomial models with a logit link. Three alternative modeling approaches were compared: (i) an individual-level generalized linear model (GLM) including population, climate treatment, and their interaction as fixed effects; (ii) a generalized linear mixed model (GLMM) including mother identity as a random intercept to account for non-independence among seedlings from the same maternal family; and (iii) a mother-aggregated binomial GLM, in which the number of surviving and non-surviving seedlings was summarized per mother, population, and climate treatment using a cbind(alive, dead) formulation. Model comparison based on Akaike’s Information Criterion (AIC) indicated that the mother-aggregated binomial GLM provided the best-supported and most parsimonious description of the data, while avoiding pseudo-replication (supplementary table S2b). This model was therefore retained for inference. Post hoc pairwise comparisons were conducted using estimated marginal means (EMMs) with Tukey adjustment for multiple testing. Statistical significance was assessed on the link (logit) scale, while effect sizes and predicted survival probabilities with 95% confidence intervals were reported on the response (probability) scale for biological interpretation. All analyses were performed in R using the packages lme4 (Bates et al. 2015), emmeans (Lenth 2024), dplyr (Wickham et al. 2023), ggplot2 (Wickham 2016), and performance (Lüdecke et al. 2021). Note that the df = Inf values reported in pairwise Tukey comparisons (supplementary table S2e) are expected for binomial GLMs. The emmeans package uses asymptotic z-tests for these models, for which denominator degrees of freedom are treated as infinite. This does not indicate an error; it is the standard behavior for GLM objects in R.

#### Trait correlations and variations

Because the experiment was conducted over two years, we tested for a year effect on phenotypic trait variation with a linear mixed-effects model implemented in the R package lme4 v1.1-35.5 (Bates et al. 2015); R Core Team, 2017):

$Y_{ij} = \mu+ {Year}_{i} +\varepsilon_{ij}$ (Eqn 1)

where *Yᵢⱼ* is the phenotypic trait measured in year *i* (AGR leaf, AGR height, RGR leaf, RGR height, Chl, Flav, NBI, C/N stem, C/N root, C/N leaves, root/shoot, SLA, leaf area), *μ* is the overall mean, and *εᵢⱼ* is the residual error.

To correct for this year effect, the best linear unbiased predictor (BLUP) values were estimated using a random-effects model:

$Y_{ij} \sim\mu+ \left( \frac{1}{Year} \right)_{i} +\varepsilon_{ij}$ (Eqn 2)

The estimated random effect of year was then subtracted from each individual's trait value to obtain a corrected value:

$Y_{ij} adjusted = Y_{ij} - coefficient\left( \frac{1}{Year} \right)_{i}j$ (Eqn 3)

Correlations and redundancy among corrected phenotypic traits were estimated using the corrR v0.4.5 function. Principal component analysis (PCA) was performed using the R package FactoMineR v2.11 (Lê et al. 2008) to assess how the 12 continuous traits (excluding survival, which was binary) contributed to overall phenotypic variation. Year-adjusted phenotypic values were subsequently used in all models testing population, climate treatment, and their interaction (Eqn 4). Experimental year was therefore not included again as an additional term in Eqn 4.

#### Statistical models to assess the role of population and climate treatment on phenotypic traits

To test for local adaptation across climate treatments, we used linear mixed-effects models. For each trait, a linear mixed-effects model was first fitted using the lme4 package in R (Bates et al. 2015):

${Yadjust}_{ijkl}=\mu+{Population}_{i}+{Condition}_{j}+\left( Population * Condition \right)_{ij}+{(1|hybrid-index)k +(1|Mother)}_{l} +\varepsilon_{ijkl}$ (Eqn 4)

where i, j, k, and l denote population, climate condition, hybrid index, and mother identity, respectively. Y*adjust*_ijkl_ represents one of the 12 growth, photosynthetic, stoichiometric, and morphological traits, after correction for experimental-year effects (Eqns 1–3). Mother identity and Hybrid index (*Pdom*) were included as random effects to account for non-independence among seedlings originating from the same maternal tree or variation in crop-to-wild admixture, respectively. The residuals $\varepsilon_{ijkl}$ were assumed to be normally distributed.

Heteroscedasticity was tested by comparing models allowing for unequal residual variance across years (2019 *vs.* 2021) with homoscedastic models using analysis of variance (ANOVA). Model selection was based on the lowest AIC, and models were simplified by removing interaction and main effects that the AIC did not support. Residual normality was assessed using the Shapiro-Wilk test and Q-Q plots (EnvStats package; (Millard 2013)). F-tests were used to determine the statistical significance of fixed effects.

Models were fitted across the whole dataset (five populations) and separately within each species (*M. domestica*, *M. orientalis*, and *M. sylvestris*). Because multiple *M. sylvestris* populations were available, explicit tests for local adaptation (population × condition) were performed within this species. Down-sampling was applied when necessary to account for unequal sample sizes across traits (supplementary tables S1 and S2); conclusions remained robust after these adjustments.

To further quantify population-specific plasticity and identify traits showing significant shifts across climate treatments, we applied random-slope regression mixed models (RRMMs) separately for each trait and population (Gómez et al. 2025). These models took the form:

$Y_{jlm}=\mu+{Condition}_{j}+\left( 1|{Mother}_{l} \right)+\varepsilon_{jlm}$(Eqn 5)

Each model was run separately within each population, with climate treatment (France, Armenia, Denmark, and Romania) included as a fixed effect. Maternal identity was treated as a random effect to account for the underlying genetic structure among seedlings. Models were fitted using the *lmerTest* package in R, allowing extraction of *p*-values for the fixed-effect climate treatment. For each trait × population combination, the statistical significance of climate treatment-induced plasticity was assessed based on these *p*-values. To characterize the direction and magnitude of plastic responses, estimated marginal means (EMMs) were calculated for each climate treatment using the *emmeans* package (Lenth 2024), and Tukey-adjusted post hoc tests were performed to identify which climate treatments differed. These RRMMs provided a detailed view of how trait plasticity varies by population, and whether domesticated or wild genotypes differ in their responsiveness to environmental conditions.

**RNA extraction, sequencing, and differential gene expression analysis**

Leaves from five seedlings per population (*M. sylvestris* from France, Romania, and Denmark; *M. orientalis* from Armenia; and *M. domestica*) were sampled under each of the four climate treatments (Armenia, Denmark, France, and Romania), for a total of 100 seedlings selected for RNA-seq (supplementary tables S1 and S3). For RNA-seq, seedlings were selected from different maternal families whenever possible to minimize relatedness among transcriptomic samples. Mother identity for each RNA-seq individual is reported in supplementary table S3. These seedlings were chosen to overlap as much as possible with those used for carbon-related trait measurements. On the last day of the experiment, half a leaf (at the 7th developmental stage) was flash-frozen in liquid nitrogen and stored at −80°C until RNA extraction. Because of their high phenolic content, leaf samples were ground using a QIAGEN TissueLyser II with PVP40 added. Total RNA was extracted with a NucleoSpin RNA Plant kit (Macherey-Nagel, Germany), following the manufacturer’s protocol. RNA purity was assessed using a NanoDrop® (Thermo Fisher) spectrophotometer, and samples with 1.8 < A260/A230 < 2.2 were considered high-quality. RNA-seq libraries were prepared by Novogene with mRNA enrichment using the Novogene NGS Stranded RNA Library Prep Set (PT044, internal kit) and sequenced on a NovaSeq 6000 instrument as paired-end 150-bp reads. The RNA-seq subset was not restricted to individuals with *Pdom* = 0 because cultivated ancestry occurs naturally within contemporary *M. sylvestris* populations, and restricting the dataset to non-admixed individuals would have substantially reduced replication across population × climate-treatment combinations. *Pdom* was recorded for each RNA-seq individual and is reported in supplementary table S1, but was not included in the DiCoExpress differential-expression model. To assess whether cultivated ancestry disproportionately influenced transcriptomic variation, we examined individuals with elevated Pdom in both a genetic network reconstructed from RNA-seq-derived SNPs and a PCA of normalized gene-expression profiles (supplementary figs. S1 and S2). In both analyses, individuals with elevated Pdom remained broadly associated with their predefined population groups and did not form a distinct cluster, indicating that cultivated ancestry was not the primary driver of the transcriptomic structure.

Raw RNA-seq reads were first quality-checked with FastQC v0.11.2 (Andrews 2010), and adapters were removed using Fastp v0.20.0 and SortMeRNA v4.3.2 (Kopylova et al. 2012; Chen et al. 2018). Cleaned reads were mapped to the *M. domestica* GDDH13 reference genome version 1.1 (Daccord et al. 2017) using STAR v2.7.10b (Dobin et al. 2013), and gene expression was quantified with Salmon v1.10.0 (Patro et al. 2017) in alignment-based mode.

Differential gene expression analysis was conducted using the DiCoExpress pipeline (Lambert et al. 2020). Differential gene expression analyses were performed twice: first, including all four climate treatments; and second, after excluding the Armenian treatment, which was sampled at a different developmental stage due to experimental constraints. This second analysis removed the only treatment affected by differences in sampling time while retaining the three climate treatments sampled under a common experimental schedule. The DiCoExpress pipeline was run using a false discovery rate (FDR) threshold of 0.05, trimmed mean of M-values (TMM) normalization (Robinson and Oshlack 2010), and a filtering threshold of counts per million (CPM) > 5. Because samples from the two experimental years also corresponded to different climate treatments and sequencing batches, count matrices were generated separately for each year and subsequently merged for differential expression analysis. Only genes showing an absolute log2 fold change > 1 (equivalent to a fold change > 2) were considered differentially expressed. Generalized linear models (GLMs) were then used to test the effects of population, climate treatment, and their interaction on gene expression.

To examine gene expression variation, normalized gene counts were modeled as a function of population, climate treatment, and their interaction using a general linear model:

${GE}_{ijk} \sim\mu+{Population}_{i}+{Condition}_{j}+{(Population * Condition)}_{ij}+\varepsilon_{ijk}$ (Eqn 6)

Where ${GE}_{ijk}$ is the normalized expression level of a given gene for a seedling *k* from population *i* under condition *j*, μ is the overall mean, and $\varepsilon_{ijk}$ *is* the residual. The interaction term captures potential genotype-by-environment (G × E) interactions. Differential expression was assessed from this full factorial model rather than from independently fitted pairwise models. Multiple testing across genes was controlled using the Benjamini–Hochberg false discovery rate (FDR) procedure.

Differential expression analyses were initially performed across all four climate treatments. Because the Armenian treatment differed from the other treatments in sampling time and showed enrichment for circadian-related expression, we additionally performed a conservative analysis restricted to the Danish, French, and Romanian treatments, which shared a common sampling schedule. In this three-treatment analysis, three pairwise climate-treatment contrasts were available within each population (Denmark–France, Denmark–Romania, and France–Romania). For each population, genes significant in at least two of these three contrasts (FDR < 0.05 and |log_2 FC| > 1) were retained as climate-treatment DEGs. Core climate-treatment DEGs were then defined as genes that met this criterion across all five populations. This analysis identified 394 core climate-treatment DEGs, of which 344 (87%) were also identified as core DEGs in the four-treatment analysis. We therefore focused subsequent biological interpretation on this conservative three-treatment core set to minimize potential confounding by sampling time.

To assess whether the functional interpretation of the core climate response was sensitive to the stringent requirement that genes respond across all five populations, we additionally applied a relaxed criterion retaining genes showing a significant climate response in at least two of the five populations. This relaxed definition identified 2,372 climate-responsive DEGs. Despite the substantially larger gene set, GO enrichment recovered the same major biological themes as the stringent core set, particularly processes related to nutrient and phosphate responses, cell-wall organization and biosynthesis, and carbohydrate and polysaccharide metabolism (supplementary fig. S7). Thus, the main functional interpretation of the shared transcriptional response to climate treatment was robust to the threshold used to define core climate-treatment DEGs.

For population DEGs, analyses were likewise based on the three climate treatments, which shared a common sampling schedule. For each of the 10 possible pairwise comparisons among the five populations, expression was compared separately within the Danish, French, and Romanian climate treatments. A gene was retained as differentially expressed between a given pair of populations when it was significant in at least two of these three climate-treatment contrasts and showed the same direction of expression change across the retained contrasts. For comparison, in the initial four-treatment analysis, six pairwise climate-treatment contrasts were available within each population, and genes significant in at least four of the six contrasts were retained as climate-treatment DEGs for that population. Core climate-treatment DEGs were defined analogously as genes meeting this criterion across all five populations.

The overlap and specificity of DEGs across comparisons were visualized using UpSet plots generated with the UpSetR package v1.4.0 (Conway et al. 2017). Population DEGs were further categorized as population-specific when a focal population was consistently differentially expressed relative to all other populations, while no significant expression differences were detected among the remaining populations. Climate-treatment DEGs were subsequently classified according to their distribution across populations as core climate-treatment DEGs (shared across all populations), species-specific DEGs (specific to *M. domestica*, *M. orientalis*, or *M. sylvestris*), population-specific DEGs within *M. sylvestris*, and wild-specific DEGs (detected only in wild species and absent from *M. domestica*).

To identify candidate expression patterns consistent with local adaptation within *M. sylvestris* (Fig. 5b), we subsequently examined climate-treatment contrasts extracted from the fitted differential-expression model separately for the Danish, French, and Romanian populations. For each population, candidate genes were defined as those showing a consistent direction of differential expression in the population’s home climate treatment relative to the other climate treatments (FDR < 0.05 and |log2FC| > 1). These climate-treatment-aligned candidates therefore differ from the population-specific DEGs shown in Fig. 3d: Fig. 3d captures expression divergence among populations, whereas Fig. 5b captures population-specific expression responses aligned with the native climate treatment.

Finally, the sets of DEGs were compared to previously identified genes under positive or balancing selection in wild and cultivated *Malus* populations (Chen et al. 2026), to evaluate whether expression patterns were associated with genomic signatures of selection.

**Assessment of cultivated ancestry in RNA-seq individuals**

To assess whether cultivated ancestry (*Pdom*) influenced the transcriptomic results, we performed two complementary analyses on the 100 RNA-seq individuals. First, genome-wide SNPs called from the RNA-seq data (see section 'Genetic divergence between populations' below) were used to reconstruct genetic relationships among the RNA-seq individuals using a bitwise genetic distance matrix computed with the poppr R package [48]. A neighbor-net split network was constructed from this distance matrix using the phangorn R package (Schliep 2011) and visualized, with nodes color-coded by hybrid index (*Pdom*), where darker shades indicate higher proportions of *M. domestica* ancestry (supplementary fig. S2a). This network was used to verify that individuals clustered primarily according to their predefined population group and that individuals with elevated *Pdom* did not form a separate admixed cluster disrupting population structure.

Second, to examine whether hybrid ancestry influenced gene expression profiles independently of population assignment, a principal component analysis (PCA) was performed on TMM-normalised gene expression values (log2-CPM) for all 96 RNA-seq individuals using the FactoMineR R package (Lê et al. 2008). Individuals were color-coded by *Pdom* in the PCA biplot (supplementary fig. S2b). Individuals with higher *Pdom* values were examined for displacement from their population group in expression space. The absence of systematic displacement of high-Pdom individuals from their population cluster in either the genetic network or the expression PCA was interpreted as evidence that cultivated ancestry did not disproportionately drive the transcriptomic patterns reported in this study.

#### Gene ontology (GO) enrichment analysis

To functionally characterize the DEGs, gene ontology (GO) term enrichment analysis was performed using the R package clusterProfiler v4.6.0 (Xu et al. 2024). The reference database (OrgDb) was generated in-house using AnnotationForge (Carlson and Pagès 2024) based on annotations of the GDDH13 reference genome of *M. domestica*. Enrichment analysis was conducted using the Benjamini-Hochberg (Benjamini and Hochberg 1995) correction for multiple testing, with both adjusted *p*-value and *q*-value thresholds set at 0.05. The background genes comprised the expressed genes retained for differential gene expression analysis (Wijesooriya et al. 2022), thereby enabling interpretation of enrichment in the context of active transcription under the tested conditions.

Relationships among significantly enriched GO terms were visualized as a semantic similarity network using the enrichplot framework from clusterProfiler. Pairwise semantic similarity between GO terms was calculated using the Wang method. GO terms were connected by an edge when their semantic similarity exceeded 0.7; terms below this threshold were left unconnected. Edge width was proportional to semantic similarity, and node size represented the number of DEGs associated with each GO term.

#### Linking phenotypic and gene expression variation

To explore the relationship between gene expression and phenotypic variation, a weighted gene co-expression network analysis (WGCNA) was performed using the R package WGCNA v1.7.3 (Langfelder et al. 2010). WGCNA constructs correlation networks to identify modules (i.e., clusters) of co-expressed genes and to relate these modules to external traits. A signed network was built using a soft-thresholding power of 12, a minimum module size of 30 genes, and a maximum block size of 10,000. Modules were summarized by their eigengene expression profiles and tested for correlation with phenotypic traits measured across seedlings.

To link co-expression patterns with the DEG analysis, the overlap between core-climate DEGs and WGCNA modules was assessed by performing enrichment analysis using Fisher’s exact tests. For each gene module, a 2 × 2 contingency table was constructed by cross-classifying genes according to (i) membership in the tested module (inside vs. outside the module) and (ii) core climate-treatment DEG status (core DEG vs. non-core DEG). To account for multiple hypothesis testing across all categories and directions, p-values were adjusted using the Benjamini-Hochberg False Discovery Rate (FDR) method and a threshold of adjusted p-value < 0.05 was used.

GO enrichment analysis was performed within each DEG-enriched module to identify potential overrepresentation of specific biological processes or molecular functions, providing insights into the functional relevance of gene expression plasticity.

**Genetic divergence between populations**

To assess whether genes differentially expressed in response to climatic variation were also associated with elevated population differentiation, the overlap between them was examined. Pairwise genetic divergence between *M. domestica*, *M. orientalis*, and each *M. sylvestris* population was estimated using *D_XY_*, calculated per exon with Pixy (Korunes and Samuk 2021). Gene-level *D_XY_* values were obtained by aggregating the Pixy metrics count_diff and count_comparison across all exons within a gene.

The SNP dataset used was generated using the following pipeline (Sauvage et al. 2017): variants were called per sample from BAM files using GATK HaplotypeCaller v4.4.0.0 (Auwera and O’Connor 2020) against the GDDH13 v1.1 reference genome. Single-sample genomic VCFs (gVCFs) were generated with --emit-ref-confidence GVCF and a minimum confidence threshold of 20. Individual gVCFs were combined into a multi-sample cohort using GATK CombineGVCFs, followed by joint genotyping with GenotypeGVCFs using the -all-sites option to retain both variant and invariant positions. Raw VCFs were filtered to retain high-quality variants in a multi-step pipeline. Sites missing critical annotations (F_MISSING = 1) were removed using BCFtools v1.14 (Danecek et al. 2021). Remaining sites were annotated and flagged using GATK VariantFiltration based on recommended hard-filtering parameters: SNPs with QUAL < 30, QD < 2.0, ReadPosRankSum < -8.0, FS > 60.0, SOR > 3.0, or MQ < 40.0 were flagged, with InDels/mixed variants subject to adjusted thresholds (ReadPosRankSum < −20.0, FS > 200.0, SOR > 10.0). Finally, all flagged sites were removed using GATK SelectVariants, yielding a VCF containing only high-confidence variants. SNPs were then annotated using SnpEff (Cingolani et al. 2012) and classified into synonymous and nonsynonymous variants using SnpSift. At the genotype level, calls with read depth (DP) < 4 or > 200 or genotype quality (GQ) < 20 were set to missing. Sites with >20% missing genotypes were subsequently excluded, and only biallelic SNPs were retained for downstream analyses.

*F_ST_* and *D_XY_* values among populations were computed using only synonymous SNPs and invariant sites, as required by Pixy to avoid bias. *D_XY_* values were standardized using Z-scores, and genes with Z-scores greater than 2 were considered highly divergent, corresponding to approximately the upper 5% of the genome-wide distribution—a commonly used threshold for identifying loci with elevated genetic divergence. Because SNPs were called from RNA-seq reads, *D_XY_* estimates are necessarily restricted to genomic regions with sufficient transcriptomic coverage and should therefore not be interpreted as genome-wide estimates of local genetic divergence. Accordingly, the overlap between *D_XY_* outliers and DEGs was interpreted conservatively as evidence of convergence between transcriptional and genetic differentiation at the gene level, rather than as evidence that expression differences were caused by cis-regulatory variation. This overlap was interpreted as convergence between transcriptional and genetic differentiation at the gene level, rather than as evidence that expression differences were caused by cis-regulatory variation. The RNA-seq-derived SNP dataset was used for these analyses because whole-genome sequencing was not available for the ecotron seedlings. Thus, the individuals used for *D_XY_* belonged to the ecotron RNA-seq dataset and were distinct from the 64 WGS individuals used for GEA. This approach also ensured that genetic divergence and differential expression were evaluated using the same experimental material.

**Genome-climate associations in *Malus sylvestris***

Genome–environment association (GEA) analyses were conducted in *M. sylvestris* using an independent whole-genome sequencing dataset comprising 5,524,908 filtered SNPs from 64 individuals sampled in France, Denmark, Austria, and Romania (Chen et al. 2026). GEA was conducted using the complementary Samβada and LFMM approaches while accounting for population structure. Significant GEA SNPs were assigned to nearby genes using a 2-kb proximity criterion; the detailed SNP-to-gene assignment procedure is described in Supporting Text S2. Full details of SNP filtering, population-structure analyses, environmental-variable selection, GEA models, significance thresholds, and gene assignment are provided in Supporting Text S2.

To determine whether the overlap between GEA candidate genes and climate-treatment DEGs exceeded the expected level by chance, we performed a permutation analysis with 10,000 randomizations. At each iteration, gene sets of equivalent size to the observed candidate sets were randomly sampled from the 19,533 genes eligible for both analyses, and their overlap was recorded. The observed number of overlapping genes was compared with the resulting null distribution, and an empirical P-value was calculated as the proportion of permutations that yielded an overlap greater than or equal to the observed value.

##### Effects of domestication on molecular diversity, gene expression, and deleterious mutation load

To assess how domestication has shaped patterns of gene expression and molecular diversity, different categories of expressed genes were compared: (1) DEGs between *M. domestica* and wild populations (“Dom-wild DEGs”), (2) core climate treatment-responsive DEGs shared across taxa (“core climate treatment DEGs”), and (3) all remaining expressed genes (“Background genes”).

Nucleotide diversity (*πN* and *πS*) was calculated with Pixy (Korunes and Samuk 2021), using 1,086,764 SNPs called from exons of 19,533 expressed genes detected in this study (see above for the SNP calling method). Average *πN* and *πS* values were then computed for each gene.

To compare molecular diversity between gene groups and between wild and domesticated populations, generalized linear mixed models (GLMMs) were fitted using the glmmTMB package in R (Brooks et al. 2017). Because the diversity metrics contained true zeros, we utilized a zero-inflated Gamma distribution (with a log link function) rather than log-transforming the response variables. Different parameterizations for the zero-inflation component were tested and compared using the performance package in R (Lüdecke et al. 2021) to assess model fit. Based on these comparisons, the final optimal models incorporated both gene category and population into the zero-inflation formula. The conditional components of the models were formulated as follows:

$\frac{\pi Ndomesticated}{\pi Nwild}$ ~ gene categories + (1|gene) (Eqn 7).

$\frac{\pi Sdomesticated}{\pi Swild}$ ~ gene categories + (1|gene) (Eqn 8)

$\frac{\pi N}{\pi S}$ ~ population + gene categories + (gene categories × population) + (1|gene) (Eqn 9)

The first two models assess changes in genetic diversity between domesticated and wild apples, and the third model tests for differences in the relative contribution of nonsynonymous versus synonymous diversity across populations and gene categories. Genes for which the corresponding wild-population denominator (*πNwild* in Eqn 7 or *πSwild* in Eqn 8) was zero were excluded from the ratio models to avoid undefined ratios. For the *πN*/*πS* model (Eqn 9), genes with *πS* = 0 were excluded, whereas genes with *πN* = 0 were retained and contributed to the zero-inflated component of the model.

The accumulation of deleterious mutations was examined within the same gene category using whole-genome SNPs from Chen et al., 2026 (Chen et al. 2026), in which derived deleterious alleles were already annotated using SIFT 4G (Vaser et al. 2016) and synonymous variants were extracted. To control for individual- and category-level variation in overall mutation rate, gene size, and baseline neutral diversity, deleterious mutation counts were evaluated relative to synonymous counts. For each individual, deleterious genotypes were quantified separately according to genotype state: heterozygous deleterious genotypes (*He*) and homozygous derived deleterious genotypes (*Ho*). The two genotype classes were analyzed separately because their relative abundance can differ among populations and therefore describe distinct components of deleterious variation. Values were then grouped according to gene category (Dom-wild DEGs, core climate treatment DEGs, and background genes).

Heterozygous (He) and homozygous (Ho) mutation counts were analyzed separately using binomial Generalized Linear Mixed Models (GLMMs) with a logit link function according to the following model:

cbind(deleterious, synonymous) ~ population + gene categories + (gene categories × population) + (1|genotype) (Eqn 10)

where genotype identity was included as a random intercept (1|genotype) to account for repeated measures across gene categories within individuals and to control for baseline individual-level variation.

Under this logistic model framework, the logit link models the log-odds of a variant being deleterious rather than synonymous (ln(deleterious/synonymous)). Post hoc contrasts and estimated marginal means were evaluated on the response scale using emmeans (type = "response"), yielding back-transformed probabilities (P = deleterious/ (deleterious + synonymous) that represent the modeled proportion of tracked mutations that are deleterious across populations and gene categories.

To control for unequal gene set sizes, random subsampling was performed at 10%, 20%, 50%, 80%, and 100% of each gene set, with the procedure repeated 100 times per subset. This approach allowed us to verify whether the observed differences in deleterious mutation load were robust against sampling bias.
