## Supplementary text 2 for "Conserved transcriptional plasticity, not local adaptation, dominates early climate responses across wild and cultivated apples"

**Extended methods** - **Text S2. Genomic basis of climate adaptation in *Malus sylvestris***

To investigate climate adaptation in *Malus sylvestris*, we combined genome-environment association (GEA) analysis with differential gene expression and genomic divergence metrics. This integrative approach aimed to identify genes that respond to climatic gradients at both the regulatory and sequence levels.

**Bioclimatic variables and preparation of environmental data**

The 19 standard bioclimatic variables (BIO1-BIO19) were extracted from the WorldClim2 database (Fick and Hijmans 2017) for all sampling sites. After filtering for collinearity using Pearson’s correlation coefficient (threshold *r* > 0.7), six non-redundant variables were retained for GEA: BIO1 (annual mean temperature), BIO2 (mean diurnal range), BIO3 (isothermality), BIO4 (temperature seasonality), BIO8 (mean temperature of the wettest quarter), and BIO12 (annual precipitation). These variables represent independent axes of climatic variation related to temperature and precipitation.

**Single-nucleotide polymorphism (SNP) dataset and population structure**

A filtered single-nucleotide polymorphism (SNP) dataset was used in this study, consisting of 5,524,908 SNPs from 64 *M. sylvestris* individuals sampled in France, Denmark, Austria, and Romania (Chen et al. 2026). These WGS individuals were independent of the ecotron seedlings used for RNA-seq, with no individual overlap between the two datasets. Austria, although not represented in the ecotron experiment, was included in the GEA dataset to provide broader geographic and climatic representation of natural *M. sylvestris* populations. Individuals showing admixture with *M. domestica*, *M. orientalis*, or *M. baccata* were excluded based on fastStructure (Raj et al. 2014) and ancestry thresholds (supplementary table S4). SNP filtering was performed with BCFtools, applying the following thresholds: minor allele frequency (MAF) > 0.05, missingness < 10%, and exclusion of monomorphic loci. Linkage disequilibrium (LD) pruning was performed with PLINK v1.9 using a 5-kb sliding window, step size = 1 SNP, and r^2^ threshold = 0.2 (–indep-pairwise 5kb 1 0.2), yielding a smaller set of 26,009 SNPs used for structure inference. This LD-pruned subset was used exclusively for fastStructure population-structure inference. The genotype–environment association analyses (SamBada and LFMM) were performed on the full set of 5,524,908 quality-filtered SNPs.

This dataset was used to infer genetic structure using fastStructure, testing K=2 to K=10, with 20 replicates per K (Raj et al. 2014), which estimates individual admixture proportions for a given number of clusters (*K*). Cluster assignments were visualized and summarized using CLUMPAK (Kopelman et al. 2015). The maximum membership coefficient for each individual was calculated to guide cluster assignment. Individuals with membership coefficients > 0.9 were assigned to their respective clusters, while those with lower membership coefficients were labeled as admixed. The optimal number of clusters (*K*) was selected by evaluating the cross-validation error across runs and retaining the *K* value with the lowest mean error. Principal component analysis (PCA) was also performed to visualize clustering patterns in a reduced-dimensional space using the dudi function.pca function in the R package ade4 (Dray and Dufour 2007).

FastStructure revealed a clear spatial pattern of population genetic structure in European *M. sylvestris*. From K = 2 to K = 6, only two major genetic clusters were distinguishable. However, at K = 7, a third cluster emerged, separating the western populations (France and Denmark) from the Austrian and Romanian populations. Following the rationale of Puechmaille (Puechmaille 2016), we retained K = 7 as the best-tested fastStructure model, in which the additional ancestry component was represented by non-admixed individuals rather than exclusively by admixed ancestry profiles. Importantly, K = 7 denotes the complexity of the fitted fastStructure model and should not be interpreted as evidence for seven distinct genetic groups; the resulting ancestry profiles instead support three major genetic clusters. Accordingly, three latent factors were used in the subsequent LFMM analyses to account for these three major components of population structure.

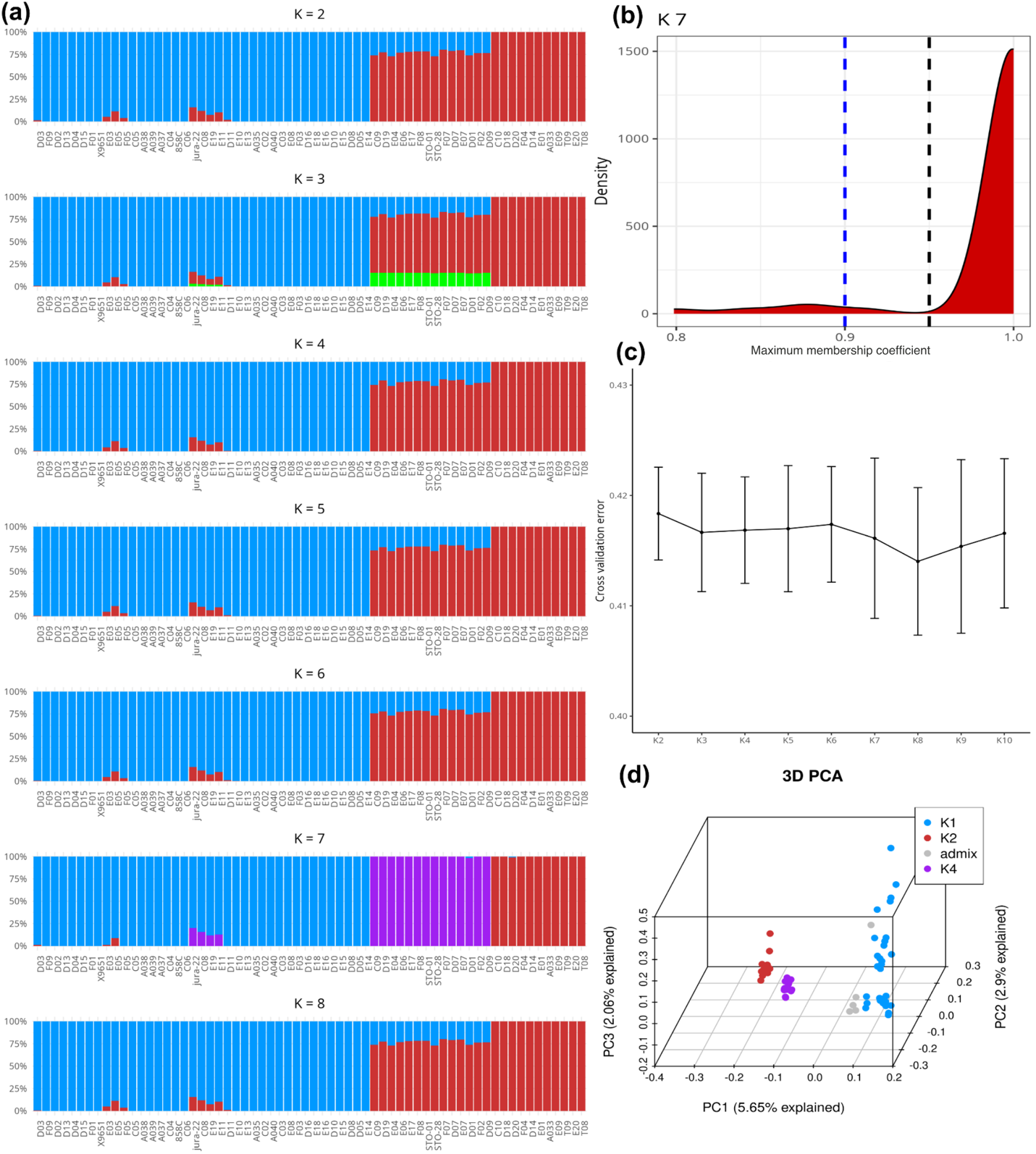

**Text S2, Fig. 1**: Population structure of 64 individuals of *M. sylvestris* from France, Denmark, Austria, and Romania. (a) Results of fastStructure analysis with *K* = 2–8 showing population assignment. (b) Distribution of the maximum ancestry membership coefficient per individual at K = 7. For each individual, the largest ancestry coefficient among the seven inferred components was retained. Dashed lines indicate assignment thresholds of 0.90 (blue) and 0.95 (black). (c) Cross-validation error across fastStructure models from K = 2 to K = 10. Points indicate mean cross-validation error across 20 independent runs, and error bars indicate ±1 SD. (d) PCA plot showing the first three principal components (PCs) at *K* = 7. Blue circles: western populations; red circles: Romanian individuals; purple circles: Austrian individuals.

**GEA methods: LFMM and Samβada**

GEA analysis was performed using two complementary methods that correct for structure:

- **LFMM** (Frichot et al. 2013) uses latent factors to capture hidden population structure. We used the optimal number of latent factors (*K*) inferred by fastStructure above. LFMM was run independently for each of the six bioclimatic variables. SNPs with *q*-values < 0.01 were considered significant.
- **Samβada** (Duruz et al. 2019)**:** We used the multivariate logistic regression option, correcting for structure by including ancestry coefficients. Samβada models were run in parallel for each environmental variable. SNPs were retained at *p* < 0.001, adjusted using the Bonferroni correction.

Environmental data were extracted from the WorldClim2 database, specifically the 19 standard bioclimatic variables (BIO1–BIO19). Following collinearity filtering (|r| > 0.7), six non-redundant bioclimatic variables were retained for subsequent GEA analyses.

**Gene-level assignment and overlap with differentially expressed genes (DEGs)**

Significant SNPs identified by both LFMM and Samβada were mapped to nearby gene models using the *M. domestica* GDDH13 v1.1 annotation (Daccord et al. 2017). SNPs located within gene bodies or within 2 kb upstream or downstream of gene boundaries were assigned to the corresponding genes.

These gene lists were compared with genes identified as differentially expressed in response to climate treatment (core climate treatment DEGs) or between populations (population DEGs), based on RNA-seq data (see Materials and Methods, main text). For each bioclimatic variable, we assessed the overlap between DEGs and GEA candidates to evaluate convergence between regulatory plasticity and genetic divergence.

**Local adaptation signals and overlap with *D_XY_* outliers**

To identify genes potentially involved in local adaptation, we used the expression profiles of DEGs to isolate genes consistently upregulated or downregulated in the home-climate treatment of each *M. sylvestris* population (relative to other populations). These gene lists were compared to genome-wide *D_XY_* outliers, calculated using Pixy (Korunes and Samuk 2021). *D_XY_* was initially calculated at the exon level, and gene-level *D_XY_* values were subsequently obtained by aggregating Pixy count_diff and count_comparison across all exons belonging to each gene before calculating the corresponding gene-level ratio. Z-scores > 2 were used to define highly divergent genes. The SNP dataset and detailed *D_XY_* calculation procedure are described in Supporting Text S1. Because these SNPs were derived from RNA-seq reads, *D_XY_* estimates are restricted to regions with sufficient transcriptomic coverage; this limitation is discussed in Supporting Text S1.

This analysis allowed us to identify a subset of candidate genes that exhibit both expression patterns consistent with local adaptation and elevated sequence divergence, providing robust evidence of climate-driven evolutionary responses.
